# Whole-body Bacteriophage Distribution Characterized by a Physiologically based Pharmacokinetic Model

**DOI:** 10.1101/2025.02.06.636931

**Authors:** Arne Echterhof, Tejas Dharmaraj, Patrick Blankenberg, Bobby Targ, Paul L. Bollyky, Nicholas M. Smith, Francis Blankenberg

## Abstract

In 2019 there were over 2.8 million cases of antibiotic-resistant bacterial infection in the US with gram negative organisms having up to a 6% rate of mortality. Bacteriophage (phage) therapy holds great promise to treat such infections. However, the biologic features which influence the pharmacokinetics (PK) of phage have been difficult to characterize due to a lack of standardized protocols of phage purification, tissue assay, and labeling. Here we present robust methods for ultrapure phage preparation as well as non-destructive highly stable attachment of radio-iodide to phage using the well described Sulfo-SHPP linker. We purified and radiolabeled the phage strains, PAML-31-1, OMKO1, and Luz24 lytic to drug-resistant Pseudomonas aeruginosa for biodistribution assay in normal young adult CD-1 mice injected via penile vein. Groups of 5 mice were euthanized and tissues/organs removed for weighing and scintillation well counting of I-125 activity at 30 min, 1h, 2h, 4h, 8h, and 24h. A physiologically based PK (PBPK) model was then constructed focusing on compartments describing blood, lung, muscle, bone, liver, stomach, spleen, small intestines, large intestines, and kidney. Model permeability coefficient (PS) was estimated across all organs as being 0.0227. Tissue partition coefficients (KP) were estimated for high perfusion organs (lung and kidney) as 0.000138, GI organs (liver, spleen, and stomach) as 0.627, and all other organs as 0.220. Elimination was governed by MPS-mediated elimination (TMPS,deg) and active secretion at epithelial barriers (CLActive), which were estimated as 0.00301 h and 0.0145 L/h/kg, respectively. Monte Caro simulations showed that the rapid elimination phage in humans is expected, resulting in phage blood concentrations being lower than 10^2^ PFU/mL (limit of quantification by plaque assay) by 12 hours. As such, multi-dose regimens and continuous infusion regimens were the only strategies that allowed continuously detectible phage concentrations. Evaluation of different dose levels showed that at a maximum dose of 10^12^ PFU, phage concentrations are expected to be approximately 10^7^ PFU/g. Our physiologically based PK model of phage represents the first rigorous pre-clinical assessment of phage PK utilizing contemporary pharmacometric approaches amenable to both pre-clinical and clinical study design.

## Introduction

Antimicrobial resistance is an ongoing public health concern and has been highlighted by the United States Centers for Disease Control(1) and Prevention and the World Health Organization(2). Presently, 2.8 M individuals in the U.S. are infected by multi or extensively drug-resistant (**MDR/XDR**) bacteria every year and is expected to grow to over 10M individuals per year by 2050(3). In the nosocomial setting, XDR *Pseudomonas aeruginosa* is particularly problematic due to their higher degree of antimicrobial resistance(4–6) and all-cause mortality rates for XDR *P. aeruginosa* have exceeded 8-30%(6–8). Unfortunately, the AMR crisis has been exacerbated by the sparse development of mechanistically novel classes of antibiotics.

Bacteriophages (phages) are viruses whose host is bacteria and can be largely characterized as lysogenic or lytic, with lytic phages being the primary focus for pharmaceutical development. Lytic phages were first suggested as possible antimicrobials in the early 20^th^ century(9), but they fell out of favor for small molecule antibiotics due to the complexities with phage mechanism of action, pharmacokinetics (PK), and resistance profile(10). Outside of their use for treating resistant bacterial infections, interest in engineering phages for drug(11, 12) or gene delivery(13) has also reinforced the need to better describe phage PK to identify optimal dosing practices(14).

Phage PK is largely dictated by their structural features as nano-scale particles but is confounded by an “auto-dosing” phenomenon that results from infecting their target bacteria(15). Baseline phage PK under uninfected conditions is poorly understood but is critically important for identifying the necessary concentration of phage at the site of infection and, ultimately, the best dose to give. Phage distribution to peripheral tissues is partially dictated by their size, which ranges 30 - 200 nm. Accordingly, the PK of phages follows many of the same principles as have been uncovered for nanoparticles and liposomal drugs over the past several decades, but additional, unique mechanisms also influence phage distribution. Phage distribution is largely permeability-limited, owed to the need to diffuse into tissues. Phage elimination is presently considered to occur through the mononuclear phagocyte system (MPS), primarily, which is phagocyte-mediated and largely occurs in the liver and spleen(16). However, phage bound to bacterial debris (e.g., endotoxin) can trigger additional immune-mediated clearance, though the exact kinetics and of these processes are still being studied(17). Though these features are currently recognized as the most influential determinants of phage PK, previous reports have shown that there are likely additional pathways to phage transport that need to be better understood.

Combined, the permeability-limited distribution to tissues and cellular-based elimination pathways, which are evolutionarily conserved processes, mean that phage PK should be scalable between species. We hypothesize that a physiologically based PK modeling approach is ideally suited to describe phage PK for pre-clinical assessment and scaling for first-in-human dose selection. To-date, phage whole body distribution *in vivo* has been characterized using limited numbers of tissues (largely blood, liver, and spleen), sparse time points, and using data-driven non-compartmental or compartmental analytical methods. Implementation of a PBPK model utilizing known anatomy of key organs (e.g., lungs, kidney, stomach, small bowel, large bowel, muscle, etc.) can improve phage translation. Additionally, quantifying the influence of phage size on disposition would be a remarkable shift in the field’s understanding of phage PK and could result in improved study designs for preclinical animal models or ongoing compassionate use situations.

Constructing a PBPK model capable of achieving these goals requires quantification methods that are amenable to rich sampling schedules and can quantify phage in both tissue and blood. Radiolabeling methods have a long tradition in pharmacology studies and have been used extensively in describing the preclinical PK of complex biologics and biotic agents^2-5^.

As such, the goal of this project was to utilize ^125^I-phage to assess the whole-body distribution of phage particles and to characterize the pharmacokinetics using a first-in-class PBPK model. As a secondary endpoint, through use of 3 different phages that are lytic with respect to most known clinically isolated strains of antibiotic-resistant *Pseudomonas aeruginosa*^6^ with unique sizes (63 nm, 227 nm, and 400 nm), we will attempt to identify any statistically significant effects of phage size on phage tissue distribution, retention, and elimination. Finally, we will attempt to scale the model for prediction of phage dosing and target site penetration in humans to hypothesize clinical dosing strategies.

## Methods

### Phage Lysate Purification

The ClMmultus OH 1 mL Monolithic Hydroxyl column with 2 μm channel size (Sartorius BIA Separations, Ajdovščina, Slovenia). was installed as per manufacturer instructions to ÄKTA Pure FPLC (GE Healthcare Biosciences, Sweden) chromatography system. Clean in place protocols (CIP) for the FPLC system and for the column were performed as per the column manufacturer’s instructions. Concentrated buffer of 200 mM Tris at pH 7.5 was used to restore pH for both CIPs. Buffers were introduced via the sample pump for the system CIP, and the injection valve for the column CIP.

Phage lysates were treated with 5 U/mL Benzonase nuclease (Sigma Aldrich) overnight at 37 °C to digest free DNA and passed through a 0.22 µm PES membrane. The subsequent lysate was then diluted at a 1:1 ratio with 3.0 M KH2PO4 buffer balanced to pH 7.0. 10 column volumes (CV) of 1.5 KH2PO4 balanced to pH 7.0 were ran through the column via injection pump. Treated phage lysates were then loaded via the injection pump into the FPLC system, and a 0-100% linear gradient was performed over 20 CV, starting with 1.5 M KH2PO4 at pH 7.0, and ending with 20 mM KH2PO4 at pH 7.0. 1 mL fractionations of column output were analyzed for a peak in UV absorbance at 280 nm using FPLC integrated wavelength UV-vis detector (GE Healthcare Biosciences, Sweden), and fractionations at the top of the UV peak were pooled and passed through 0.22 µm PES membrane. Filtered product was considered purified phage. Column was then re-equilibrated with 10 CV of 1.5 M KH2PO4 before repeating system CIP and column CIPs as per manufacturer instructions.

The PAML-31-1 and OMKO1 lytic phages were prepared with the ÄKTA/Monolithic Hydroxyl column in volume of ∼ 20 mL of 1 to 5 x 10^11^ pfu/mL with LPS < 0.05 and < 0.5 EU/mL, respectively, measured using a recombinant Factor C Endotoxin Detection assay (bioMérieux, France, Endonext EndoZyme II assay) with an assay range from 0.005 to 50 EU/mL. The Luz24 phage prep however had an average of 20 EU/mL of LPS after column purification and required with sequential spin/wash cycles x10 after overnight DNase treatment of 0.22 µm filtered lysate, spiked with rh-Annexin V (NASI, 2004), 0.1 mg/mL + 2 mM CaCl2 (final concentrations) in HEPES buffer and 15 mL 100 kDa Amicon™ centrifugal filters spun 10 minutes at 5000 xg per cycle. Two additional cycles were performed with HEPES buffer to remove Annexin V/Ca^2+^ from the phage solution. The LPS level of purified of Luz24 was 1.89 EU/mL.

All phage samples demonstrated > 98.5% reduction of proteins as measured by a bicinchoninic acid (BCA) Pierce BCA Protein Assay kit (Thermo Scientific, USA). While column purification increased DNA content post-purification in most samples the elution of labeled phage on a NAP- 5 column results in removal of > 97% of nucleotides and oligonucleotides from phage samples Note there was < 30 ng/mL of DNA before radiolabeling/NAP-5 filtration as measured by the determined by a Quant-iT PicoGreen assay (Invitrogen, USA) for all three phages.

### Radio-iodination of Sulfo-SHPP and conjugation of I-125-Sulfo-SHPP to Phage

Briefly, 15 µL (0.84 to 1.15 mCi) of stock I-125 (0.84 to 1.15 mCi, NEN Radiochemicals) in 0.1 M NaOH (pH 12-14, reductant free) was added to 75 µL of buffer (0.1 M Na2HPO4 + 100 mM NaCl, pH 7.2) in a 1 mL 6 x 50 mm borosilicate glass tube. Next three doubly buffer washed Pierce™ Iodination Beads were added immediately followed by 10 µL of a freshly prepared solution of 2 - 3 mg Sulfo-SHPP (Thermo Scientific Pierce Sulfo-SHPP, sulfosuccinimidyl-3 -(4-hydroxypheynyl) propionate dissolved in 10 mL of Na2HPO4 buffer. The Sulfo-SHPP/I-125/Iodobead mixture was then reacted for 30 min at RT.

After 30 min the Sulfo-SHPP/I-125 mixture was aspirated and placed into a second fresh glass tube. The three Iodobeads remaining in the first glass tube, were then washed twice with 50 µL of HEPES conjugation buffer (20 mM HEPES, 100 mM NaCl, pH 7.4) and the wash was placed into the second test tube containing the Sulfo-SHPP/I-125 mixture and was allowed to rest for 30 min at RT to ensure completion of the oxidation/radio-iodination reaction.

For phage conjugation the Sulfo-SHPP/I-125 mixture (∼170 to 180 µL) was added to an 0.5 mL Eppendorf tube containing 4 mL of 1 to 5 x 10^11^ pfu/mL phage suspended in HEPES buffer concentrated down to a 100 to 200 uL final volume with 100 kDa Amicon™ microcentrifugal filters. The conjugation reaction was allowed to proceed for 10 minutes with frequent gentle mixing using a pipette. After 10 minutes the conjugation reaction mixture was placed into the reservoir of a NAP™-5 desalting gel column (Cytiva, Amersham) equilibriated with 1x PBS buffer (pH-7.2) and allowed to flow through. An additional volume of 1x PBS was stacked on top of the conjugation reaction mixture to equal a total flow through volume of 500 µL. Eluted solution (500 µL) was then discarded and 500 uL of 1x PBS (Fraction #1) was placed into the NAP-5 reservoir and collected followed by 250 uL of 1x PBS (Fraction #2) which was also collected. Fractions #1 and #2 were pooled with an activity of 12.2 to 17.9 uCi of conjugated SHPP-I-125-phage in a final volume 2.5 mL of 1x PBS.

### In vivo studies

Healthy, 5 to 7 week old male Swiss-Webster CD-1 mice weighing 22 to 30 grams (Charles River) one week after acclimation to housing at the VSC facility at Stanford. All mice were housed and maintained according as approved in an animal protocol reviewed by the Administrative Panel for Laboratory Animal Care at Stanford University, using procedures specified by the Institutional Animal Care and Use Committees. Housing conditions for mice consisted of a 12:12 h dark−light cycle and temperature of 22 ± 2 °C with food and water ad libitum.

50 uL of a total phage volume of 2.5 mL was bolus injected into the penile vein of mice anesthetized with ip injection of 0.1 mL of a 7:3:1 cocktail of sterile 1x PBS, Ketamine (100 mg/mL), Xylazine (20 mg/mL), respectively. Mice were divided into six groups of five mice each euthanized via CO2 inhalation after sedation with the 7:3:1 cocktail at 30 min, 1hr, 2hr, 4hr, 8hr, 24hr after injection of radiolabeled OMKO1 and Luz24 phage. For the PAML-31-1 group mice were injected 30 min, 1hr, 2hr, 4hr, 24hr and 72 hr after injection of radiolabeled phage.

Blood samples were obtained by retro-orbital bleeds with 70 uL heparinized hematocrit tubes in separate groups of sedated mice at 1 min, 5 min, 15 min or 30 minutes after tracer injection.

After euthanization mice were dissected and target organs were harvested, washed in 1x PBS, and weighed in sample tubes for scintillation well counting. Samples as well as standard tubes (1 mL of a 1% injected activity) were counted for 1 min at an energy level of 0 - 100 keV with a window of 0 to 10 keV with subtraction of background radiation. Results were expressed as %ID (injected dose) or %ID/gram of tissue.

### Pharmacokinetic Analysis

Data were analyzed as %ID/g for blood and all tissues, whereas the stomach contents were modeled as %ID. A physiologically based PK (**PBPK**) model was focusing on compartments describing blood, lung, muscle, bone, liver, stomach, spleen, small intestines, large intestines, and kidney. Additionally, compartments for skin and brain were included as hypothesis generating, but for which no data was available. A carcass compartment was used for mass balance of drug from other tissues not measured. All organs except were modeled using a permeability-limited distribution structure, and each tissue was described as:

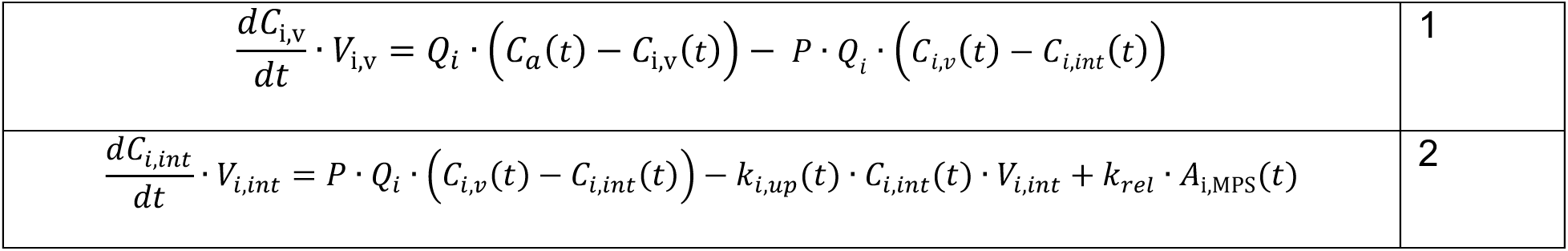

|  |  |
| --- | --- |
| $\frac{dC_{i,v}}{dt} \cdot V_{i,v} = Q_i \cdot (C_a(t) - C_{i,v}(t)) - P \cdot Q_i \cdot (C_{i,v}(t) - C_{i,int}(t))$ | 1 |
| $\frac{dC_{i,int}}{dt} \cdot V_{i,int} = P \cdot Q_i \cdot (C_{i,v}(t) - C_{i,int}(t)) - k_{i,up}(t) \cdot C_{i,int}(t) \cdot V_{i,int} + k_{rel} \cdot A_{i,MPS}(t)$ | 2 |

Where, for tissue *i*,, Cv is the phage concentration of the vascular space of the tissue, Cint is the phage concentration in the interstitial space of tissue, Vv is the vascular volume of the tissue, Vint is the volume of the interstitial space of the tissue, P is the permeability-surface coefficient, Qi is the blood flow rate of the tissue, kup is a first order rate constant governing phage uptake into the MPS compartment, krel is a first order rate constant governing phage release from the MPS compartment, and AMPS is the amount of phage in the MPS compartment of the tissue.

MPS uptake was modeled as a saturable process that was dependent on the total number of tissue resident phagocytes, maximum capacity (parameterized as %ID per 10^5^ phagocytes), and the amount of phage in the MPS compartment. This was described by the following equations:

|  |  |
| --- | --- |
| $\frac{dA_{i,res}}{dt} = k_{i,up}(t) \cdot C_{i,int}(t) \cdot V_{i,int} - k_{rel} \cdot A_{i,MPS}(t) - k_{deg} \cdot A_{i,MPS}(t)$ | 3 |
| $k_{i,up}(t) = k_{up,max} \cdot \left(1 - \frac{A_{i,MPS}(t)}{A_{i,max}}\right)$ | 4 |
| $A_{i,max} = M_i \cdot V_i \cdot \left(\frac{A_{MPS}^*}{10^5}\right)$ | 5 |
| $A_{RES}^* = \frac{A_{RES}}{Dose}$ | 6 |

Where, Ai,max is the total tissue capacity of the phage dose (%ID), Mi is the number of tissue resident phagocytes per gram of tissue (phagocytes/g), Vi is the total tissue volume (g), and Dmax is the fixed parameter of MPS carrying capacity (%ID/10^5^ phagocytes)(18). Preliminary data showing transcytosis of phage across epithelial membranes supports that a portion of phage elimination likely occurs through cell-mediated processes located at the gastrointestinal surface, hepatobiliary interface, and in the kidney. Phage degradation rate in the MPS compartment was fixed to a 2h half-life, based on preclinical cell uptake studies previously published.

Lung and kidney compartments were described using a unique partition coefficient due to each organ’s large blood flow rates. Equations describing this are described in the **supplemental materials**. Model parameters and sources are outlined in Table S1. Briefly, mouse organ weights were fixed as fractions of total weight. Mouse organ flow rates were fixed to values based on the fraction of cardiac output, which was scaled based on mouse weight.

Model was simultaneously fit to all data from each tissue, estimating a whole-body permeability coefficient (P), tissue partition coefficients (Kp), phage capacity in “MPS” sub-compartment in each tissue (Amax), and MPS transport rate (Tup). Initially, a single tissue partition coefficient was used to describe all tissues.

The structural model was estimated using SAEM in Monolix (2024R1, Lixoft) using a naïve pooled analysis (i.e., no random effects). The model was fit allowing an initial stage of parameter variability for 250 iterations to permit a full evaluation of the parameter search space. After the structural model was finalized, phage size was assessed as a possible covariate on PBPK model parameters. It was not feasible to quantify inter-animal variability because concentration data were collected as a single terminal endpoint for each animal. However, to allow for efficient testing of size as a covariate on PBPK model parameters, a single SAEM iteration was performed using the final parameter estimates for the PBPK structural model, but assuming an interindividual variability of ω^2^ = 1 on each parameter of interest. This was done to generate empirical Bayesian estimates of each parameter for each mouse to test for correlations between individual parameters and phage size. Model parameters that exhibited statistically significant correlations with phage size (>0.3) were re-estimated with a phage size covariate effect using the naïve pooled analysis strategy used to determine the structural model. The process for assessing statistically significant effects of phage or phage size effects was performed on Kp, Ps, Amax, Tup, Tdeg, and CLActive.

### Sensitivity Analysis

Local sensitivity analysis was performed to assess model changes in blood AUC in response to perturbations in Amax, CLactive, Kp,Crs, Kp,LunKid, Kp,LvrSpnSto, PS, Tdeg, and Tup. This was performed by running Monte Carlo simulations from the final model using a stepwise approach of modifying a single parameter value 5-fold greater and lower than the estimated value. Mean change in AUC was calculated as:

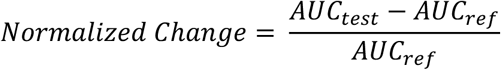

Variance-based global sensitivity analysis was performed obtaining 2000 samples of each parameter from a uniform distribution with limits that were 5-fold grater and lower than the estimated parameter value. Using the Sensitivity package(19) in R using a rank-0based estimate of the first-order sensitivity indices. Ultimately, this approach allows for efficient assessment of global sensitivity as a way to identify parameters, which can be better interrogated by future experiments.

### Interspecies Scaling

To further qualify the final model, anatomical parameters for rats and humans were used to simulate phage concentration-time profiles after administration of phage. Tissue resident macrophage concentrations were adjusted based on literature values. The full table of parameters used is located in **Supplemental Table S1.**

### External validation

The model was validated using data digitized from the literature(20, 21) using WebPlotDigitizer. Simulations were performed based on the listed dosing of phage in each study. For rats and humans, phage was assumed to administered as a rapid 5-min push.

In the seminal Kim et al study, multiple administration strategies were tested (oral), however only bolus dosing was evaluated due to unclear mechanisms of absorption associated with oral administration. Rodent studies were handled in an identical fashion. Of note, phage PK in the Kim et al. study evaluated *E. coli* phages whereas the Li et al. study evaluated *K. pneumoniae* phages. By evaluating model performance constructed on data generated on differentially sized *P. aeruginosa* phages, this external validation approach will provide information on the extrapolation potential of phage PK between phage genera. Observations versus predictions were quantitatively evaluated based on relative bias (rBias) and root mean square error (RMSE) as done previously(22).

### Clinical trial simulation

Simulation-based strategies were utilized to evaluate possible regimen structures of phage and provide a rational, data-driven basis for future trial design. Daily doses of 10^7^-10^12^ PFU were each tested with different administration schedules: single dose (x1), daily (Q24H), twice-daily (Q12H), and continuous infusion. Though phage pharmacodynamics are unquestionably important in identifying optimal regimens, standalone PK stimulations are critical to designing future *in vivo* or clinical studies that will interrogate questions related to phage-bacterial interactions.

## Results

### Whole Body Distribution of Phage and Noncompartmental Analysis

Phage concentrations in tissues varied greatly, with peak concentrations that ranged 0.6 to 15 %ID/g between all tissues analyzed. Phage concentration declined within the first 1h after administration with a half-life of ≈ 0.3 h, then exhibited a terminal half-life of ≈ 8 h. The organs with the highest concentration of all three phages included the blood, lungs, liver, spleen, and kidneys. Interestingly, a median 18.0 %ID (SD = 6.4 %ID) of phage was discovered in the lumen of the stomach 1 to 4 h after penile vein administration.

A Naïve pooled non-compartmental analysis was performed on median blood concentration values in each organ. Across all three phages, the median terminal half-life in blood was estimated as being 11.9 h, with individual half-lives for LUZ24, OMKO1, and PAML31 were calculated as 10.3, 11.9, and 12.0 h, respectively. The observed AUC0-last was 35.5 %ID*h/g after a single dose of phage across all studies. In blood, PAML31 exhibited a significantly larger AUC0-last of 195 %ID*h/g, as compared to 35.5 %ID*h/g for LUZ24 or 33.1 %ID*h/g for OMKO1.

### Physiologically Based PK Model of Phage

Final model structure is shown in **Figure 1** and model parameters are summarized in **Table 1**. Phage distribution was modeled as a permeability-limited process, where a whole-body permeability coefficient was simultaneously estimated, resulting in a final value of 0.0227. As part of the model development process, tissue partition coefficients were estimated for high flow-rate organs (liver, lung, and kidney), gastrointestinal organs whose flow collects in the hepatic portal vein (stomach, spleen, small intestines, and large intestines), and the remainder of the body (all other organs). Tissue portioning, Kp, described the likelihood of phage to be retained in each organ, and separate values were needed to describe high blood flow organs (i.e., lung and kidney) and gastrointestinal organs (i.e., Liver, spleen, and stomach). These were estimated after backward elimination, until all organs were well characterized. (**Figure 2** and **Supplemental Figures S1-S12**) First-order active clearance from surfaces in the kidney (urinary excretion) and in the liver, stomach, and intestines (gastric excretion) was a critical parameter governing elimination was estimated as 0.0145 L/h/kg. Additionally, MPS-mediated clearance was also estimated based on a degradation half-life in the MPS compartment within each tissue, which was estimated as 0.0301 h. Each tissue’s total capacity for phage was based on literature-derived values of tissue-resident phagocytes (see **Supplemental Table S1**) and a literature derived capacity of 10^3.81^ PFU/10^5^ phagocytes, based on cell uptake capacities determined *in vitro.* The uptake half-life (TMPS,up) was fixed to 0.001 with the assumption that endosomal degradation time of phage was rate-limiting. After performing a single expectation step (inter-animal variability fixed to ω = 1), correlations (>0.3) were detected between CLActive and experiments conducted using the PAML-31 phage. The inclusion of a categorical covariate to describe a PAML-31-specific reduction in CLActive resulted in a statistically significantly improved model, and the covariate was estimated as producing a 10.7-fold lower CLActive for PAML-31 (0.00134 L/h/kg). Finally, measurements obtained from stomach contents and urine provided an opportunity to characterize phage elimination through alternative pathways. Estimation of gastric transit and urine transit half-lives were done as a mechanism to empirically account for ^125^I removal from these compartments.

**Figure 1:**
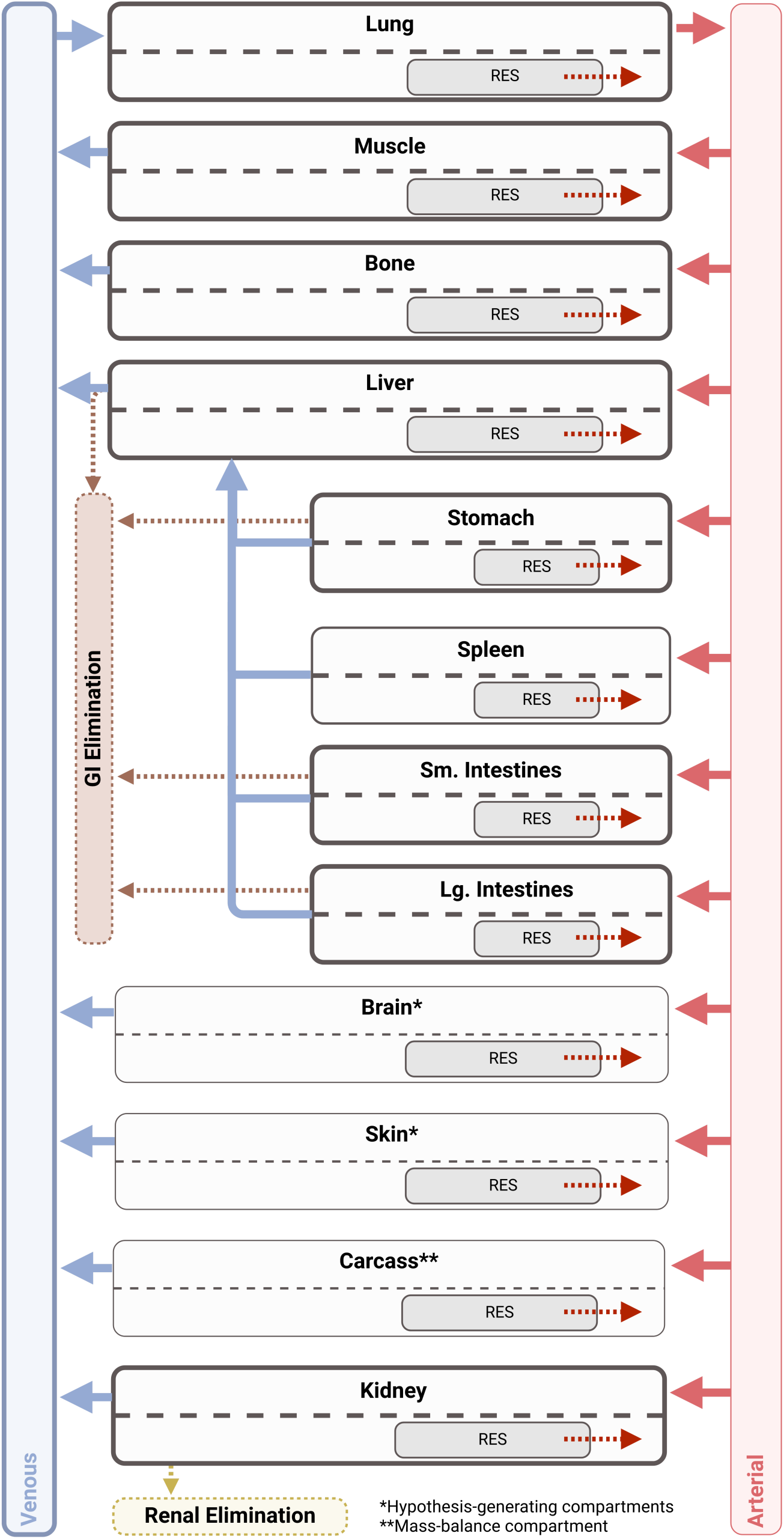
PBPK Model Structure. The final PBPK model included compartments describing the lung, muscle, bone, liver, stomach, spleen, small and large intestines, and kidney. Additional compartments were included for hypothesis generating purposes (skin and brain) along with a compartment for mass balance (carcass). Phage clearance occurred through two pathways: RES-mediated clearance in each organ and active clearance at epithelial surfaces in the kidney, stomach, and intestines. Due to unique anatomical considerations, the liver receives dual input from the hepatic artery and the Stomach, spleen, and intestines via the hepatic portal vein.

**Figure 2:**
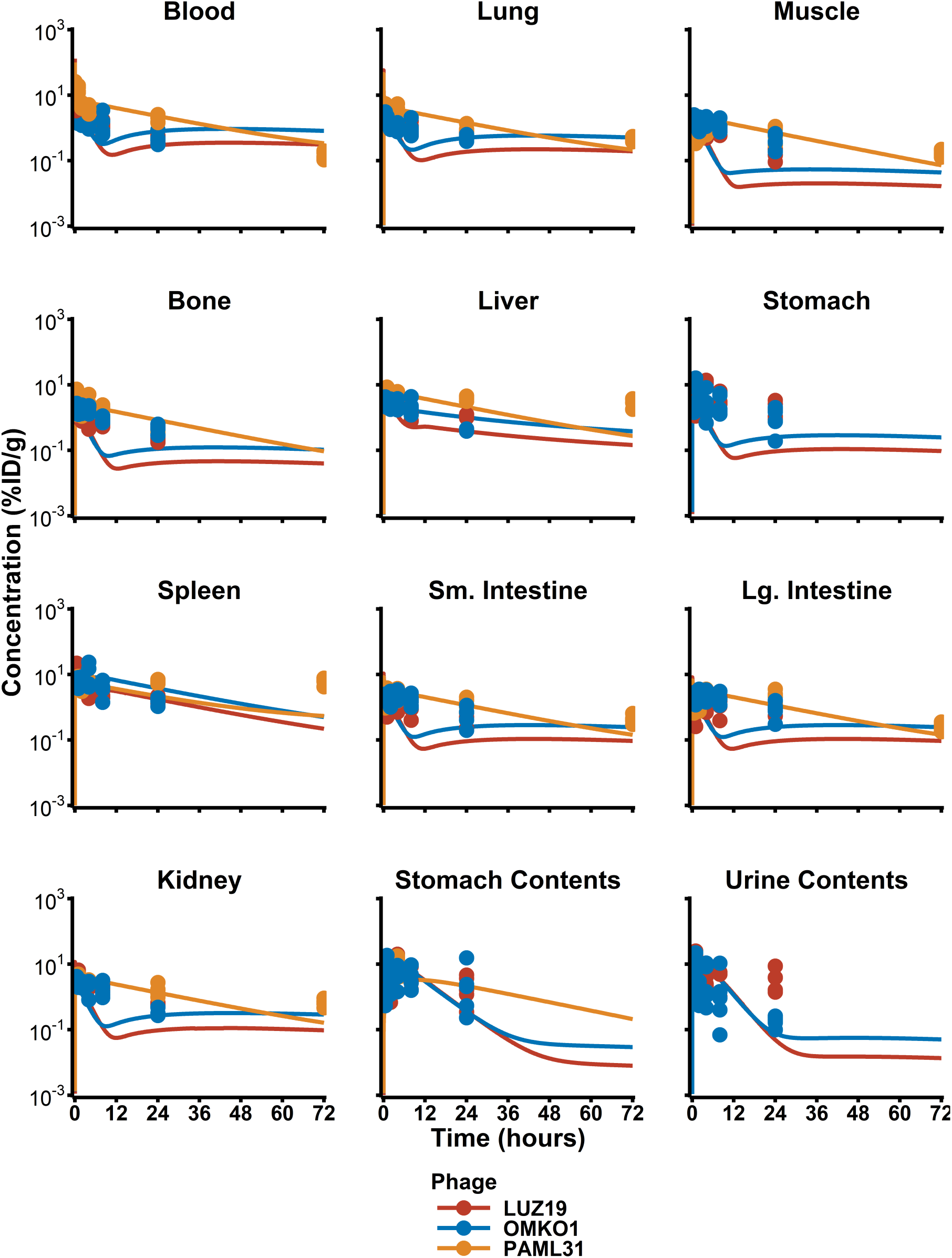
PBPK Model Post Hoc Fits. Raw data from animal replicates (dots) are presented along with PBPK model predictions (lines).

**Table 1:**
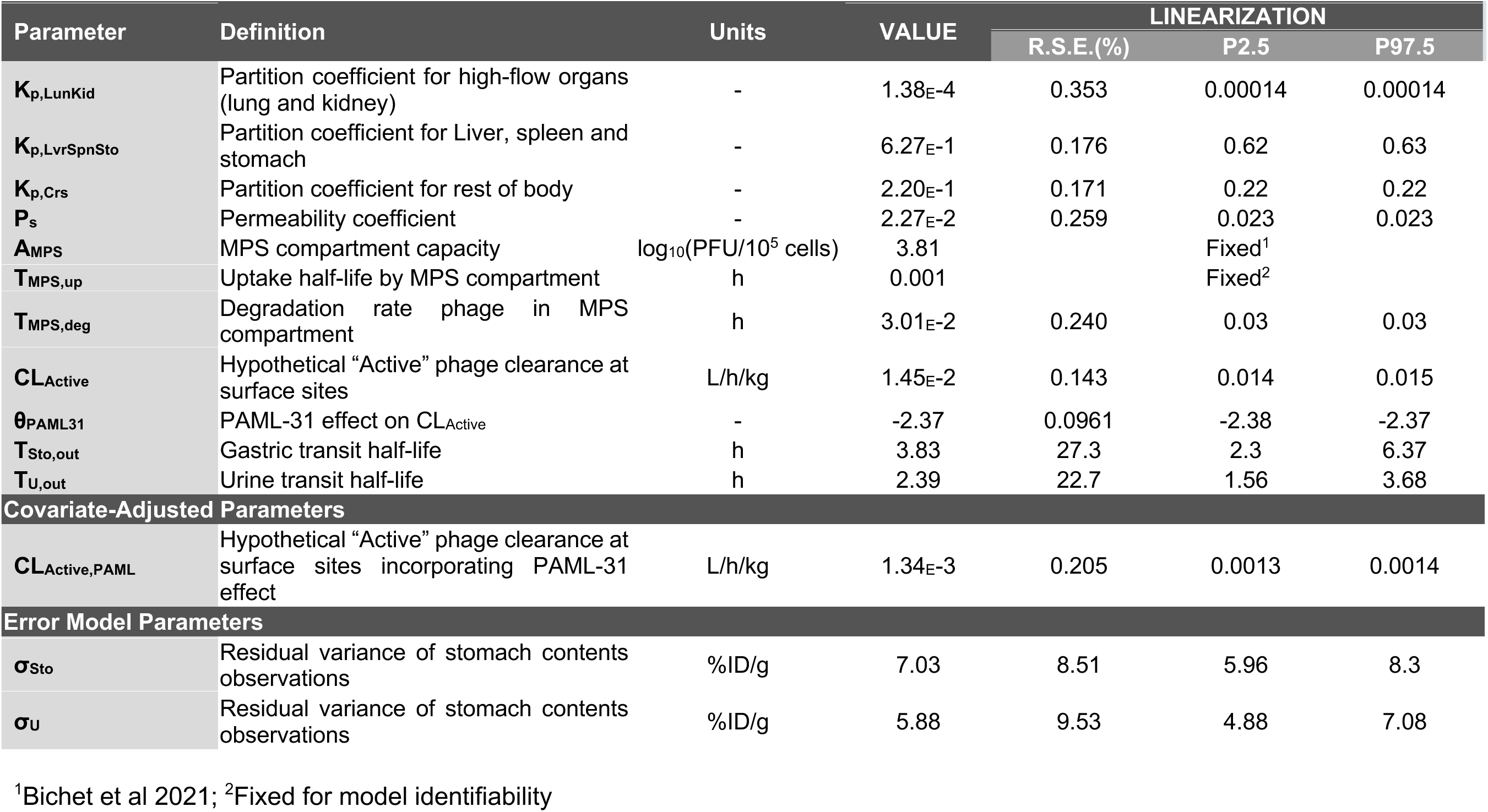
Parameter Estimates.

| Parameter | Definition | Units | VALUE | LINEARIZATION |  |  |
| --- | --- | --- | --- | --- | --- | --- |
|  |  |  |  | R.S.E.(%) | P2.5 | P97.5 |
| $K_{p,LunKid}$ | Partition coefficient for high-flow organs (lung and kidney) | - | 1.38E-4 | 0.353 | 0.00014 | 0.00014 |
| $K_{p,LvrSpnSto}$ | Partition coefficient for Liver, spleen and stomach | - | 6.27E-1 | 0.176 | 0.62 | 0.63 |
| $K_{p,Crs}$ | Partition coefficient for rest of body | - | 2.20E-1 | 0.171 | 0.22 | 0.22 |
| $P_s$ | Permeability coefficient | - | 2.27E-2 | 0.259 | 0.023 | 0.023 |
| $A_{MPS}$ | MPS compartment capacity | $\log_{10}(\text{PFU}/10^5 \text{ cells})$ | 3.81 | | Fixed<sup>1</sup> | |
| $T_{MPS,up}$ | Uptake half-life by MPS compartment | h | 0.001 | | Fixed<sup>2</sup> | |
| $T_{MPS,deg}$ | Degradation rate phage in MPS compartment | h | 3.01E-2 | 0.240 | 0.03 | 0.03 |
| $CL_{Active}$ | Hypothetical “Active” phage clearance at surface sites | L/h/kg | 1.45E-2 | 0.143 | 0.014 | 0.015 |
| $\theta_{PAML31}$ | PAML-31 effect on $CL_{Active}$ | - | -2.37 | 0.0961 | -2.38 | -2.37 |
| $T_{Sto,out}$ | Gastric transit half-life | h | 3.83 | 27.3 | 2.3 | 6.37 |
| $T_{U,out}$ | Urine transit half-life | h | 2.39 | 22.7 | 1.56 | 3.68 |
| **Covariate-Adjusted Parameters** |  |  |  |  |  |  |
| $CL_{Active,PAML}$ | Hypothetical “Active” phage clearance at surface sites incorporating PAML-31 effect | L/h/kg | 1.34E-3 | 0.205 | 0.0013 | 0.0014 |
| **Error Model Parameters** |  |  |  |  |  |  |
| $\sigma_{Sto}$ | Residual variance of stomach contents observations | %ID/g | 7.03 | 8.51 | 5.96 | 8.3 |
| $\sigma_U$ | Residual variance of stomach contents observations | %ID/g | 5.88 | 9.53 | 4.88 | 7.08 |

<sup>1</sup>Bichet et al 2021; <sup>2</sup>Fixed for model identifiability

Across all three phages in all tissues at 24h, the spleen had the longest residence time (**Figure 3**) of any organ (calculated as the ratio between the area under the first and zeroth moment curves) at 10.7 h with the liver having the second longest residence time at 8.78 h. The muscle had the shortest residence time at 6.68 h. Of note, LUZ24 experienced significantly shorter.

**Figure 3:**
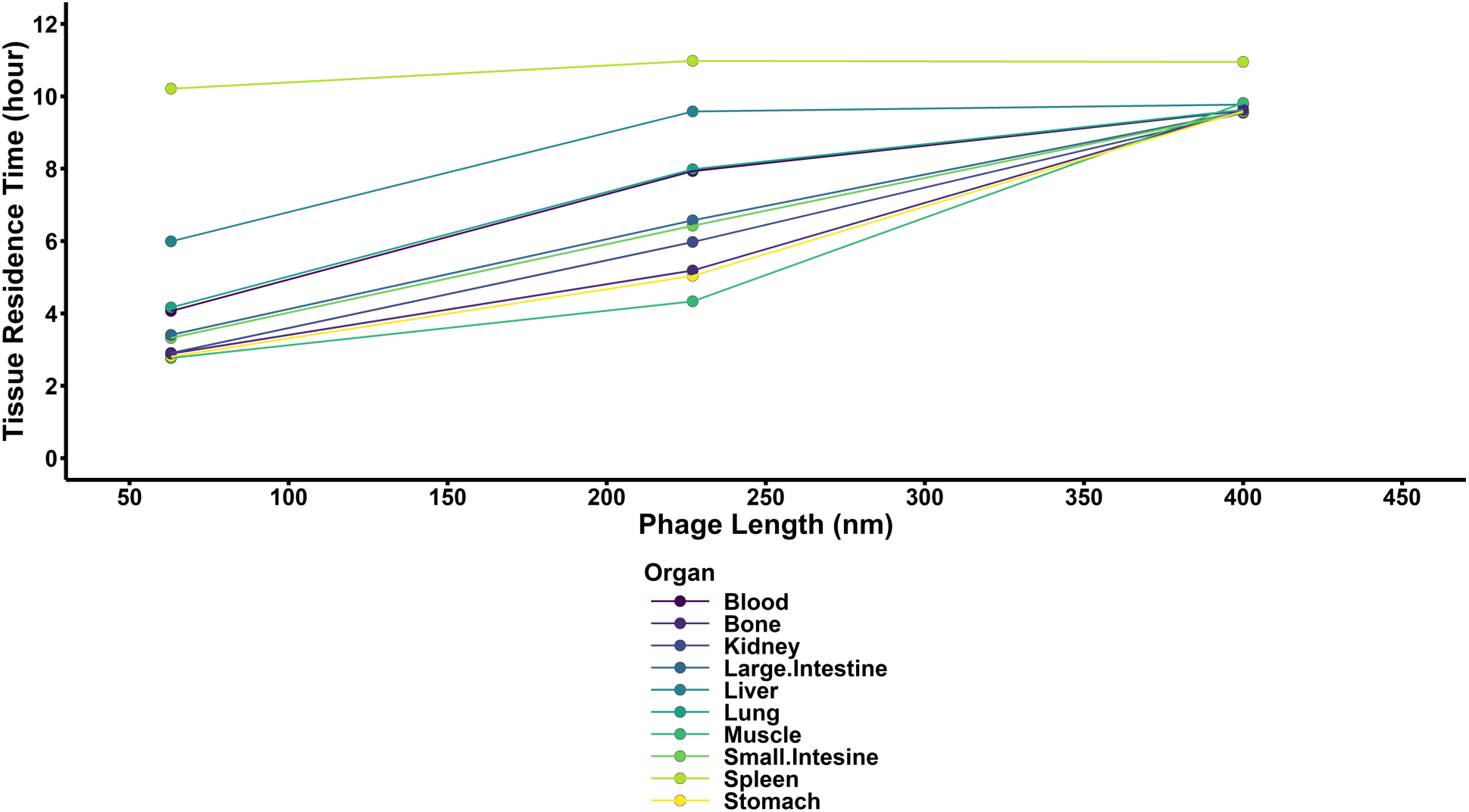
Tissue mean residence times of each phage. Tissue mean residence times were calculated numerically using the PBPK model as a ratio between the AUMC and AUC in each organ. Increased tissue residence times are associated with longer duration of phage residence in each organ and increase with size for most organs. Of note, the spleen had a more consistent residence time across all phage sizes.

### Sensitivity Analysis

The model was most sensitive (**Figure 4**) to phage capacity in the MPS (AMPS), with a Sobol Rank Index of 0.705 (Bias = 0.00126, 95%CI 0.679-0.719). After AMPS, all other parameters had absolute Sobol Rank Indices < 0.0407 (see **Supplemental Figure S13**) Next most sensitive parameters were Kp, Crs and CLActive. Of note, AMPS was sampled on a log10 scale, which provides a significantly larger search space than the other parameters.

**Figure 4:**
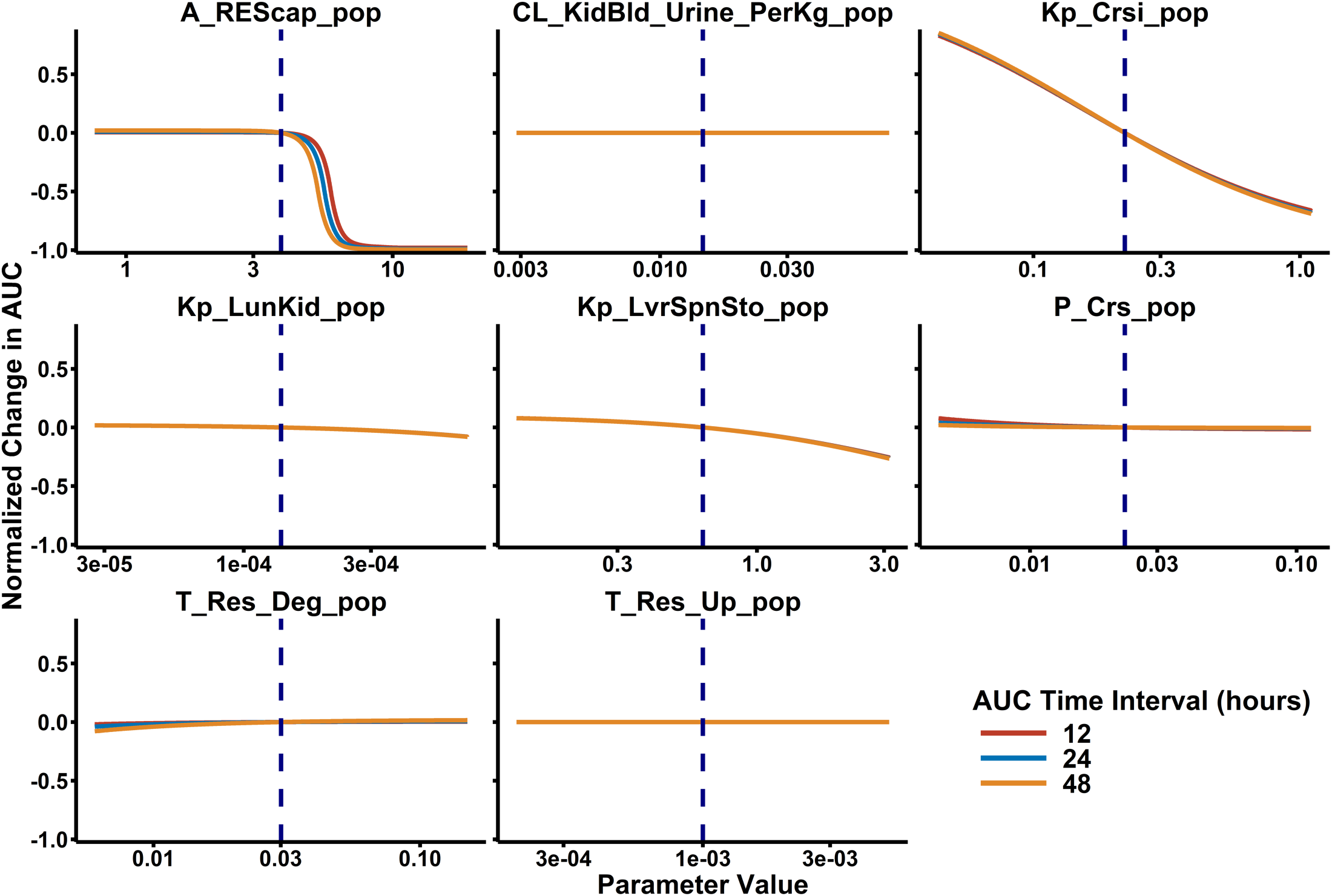
L**o**cal **Sensitivity Analysis Results.** Local sensitivity analysis was performed by sequentially modifying each parameter to a maximum difference of ±5-fold relative to the estimated value while all other parameters were fixed. In doing so, the magnitude in change in total blood exposure (i.e., area under the phage concentration-time curve) due to each PBPK model parameter can be evaluated. Sensitivity to each parameter was evaluated at 12, 24, and 48 hours after phage administration to confirm minimal time- dependent effects.

### Scaling to Humans

Human parameters are outlined in **Supplemental Table S1**, with sources. Active clearance was allometrically scaled by body mass, assuming human median mass of 70 kg. Simulation of 11 PFU intravenously every 12 hours (Q12H) showed reasonable agreement (**Figure 5**). Poor predictions primarily occurred at digitized data after 72 hours. Overall, the relative bias (rBias) in the simulation-predictions was -67.9% with a relative root mean squared error (rRMSE) of 13.1%. When only evaluating the first 72 hours of data rBias increased to -70.1% whereas rRMSE decreased to 9.71%. When attempting to simulate the PK of *Klebsiella* phages in rats, there was a significant over prediction of the highest 10^11^ PFU dose of phage, whereas the other two dose groups (10^9^ and 10^6^ PFU) were more accurately predicted.

**Figure 5:**
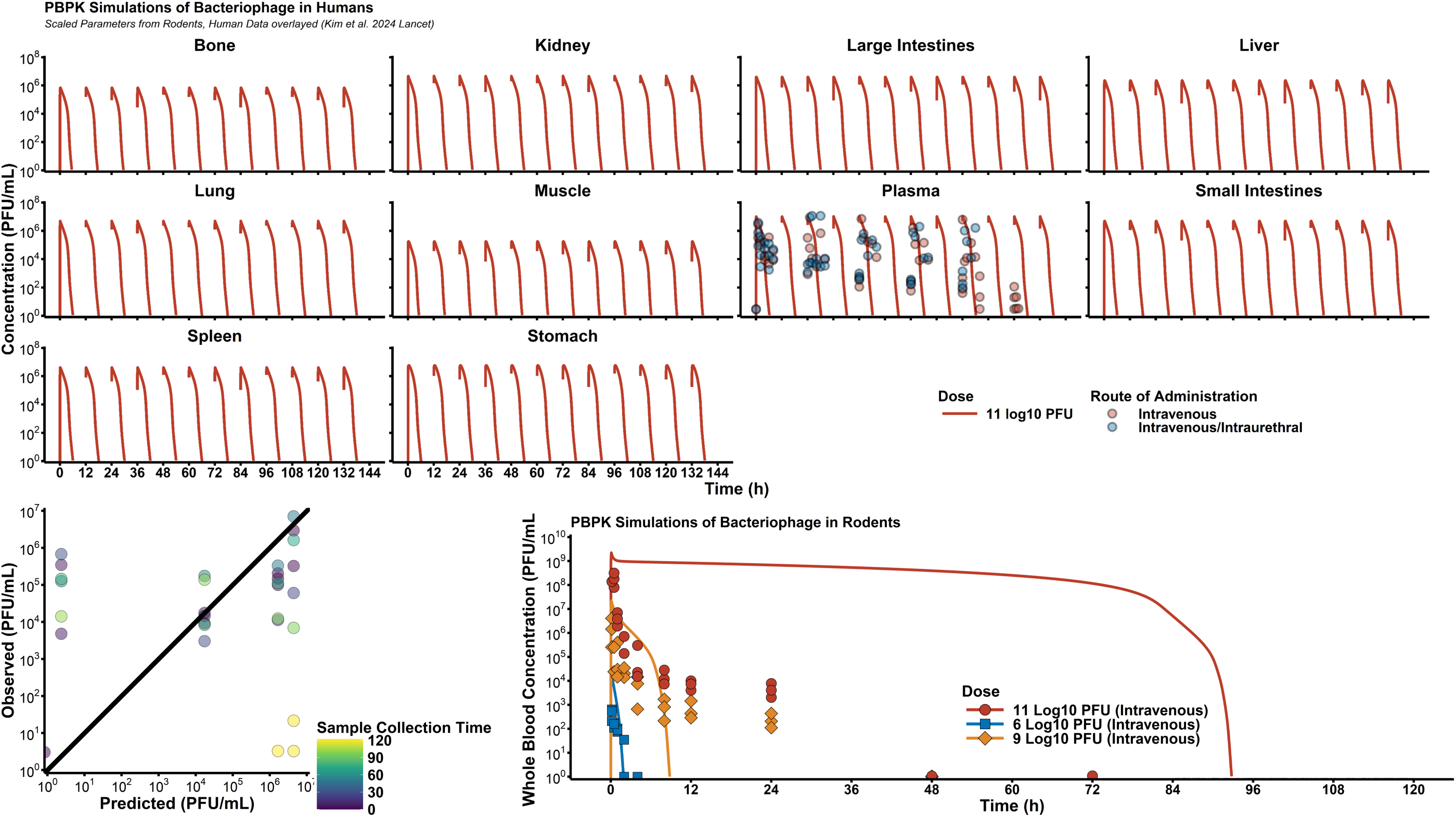
H**u**man **Simulations & Comparison to Clinical Data.** To externally validate the PBPK model, we digitized data rodent data using Pseudomonas phage øPEV20 (Lin et al, 2020, CMI) and CRISPR-enhanced *E. coli* phage LBP-EC01(Kim et al, 2024 Lancet). For human simulations, observed clinical data were compared to model predictions and relative bias (rBias) and relative root mean square error (rRMSE) were calculated to assess performance. Rodent predictions at the highest dose level was significantly biased, likely due to model misspecification of the saturable clearance processes.

Monte Caro simulations (**Figure 6**) showed that the rapid elimination phage in humans is expected, resulting in phage blood concentrations being lower than 10^2^ PFU/mL (limit of quantification by plaque assay) by 12 hours. As such, multi-dose regimens and continuous infusion regimens were the only strategies that allowed continuously detectible phage concentrations. Evaluation of different dose levels showed that at a maximum dose of 10^12^ PFU, phage concentrations are expected to be approximately 10^7^ PFU/g.

**Figure 6:**
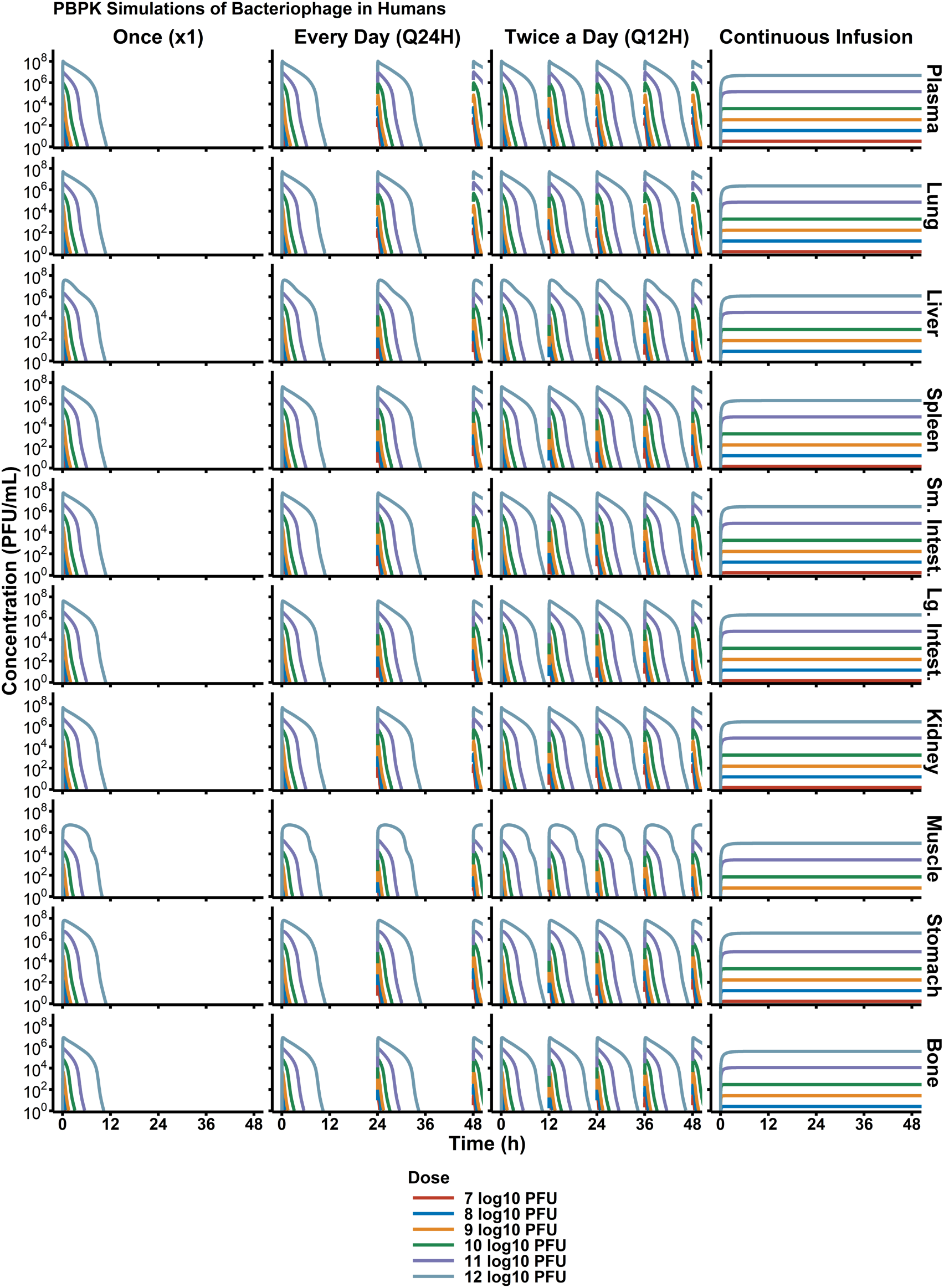
S**i**mulated **Trial of Phage PK in Humans.** To design first-in-human phage dosage regimens, simulation-based strategies are critical. Simulations of different dose levels and administration schedules (e.g., once, Q24H, Q12H) provided a robust understanding of expected phage concentrations in each organ. Overall, due to phage’s rapid elimination. With additional pharmacodynamic data, phage dosages can be then optimized to target specific, therapeutic concentrations in infection sites of interest.

## Discussion

Our physiologically based PK model of phage represents the first rigorous pre-clinical assessment of phage PK utilizing contemporary pharmacometric approaches amenable to both pre-clinical and clinical study design. Previous studies of phage PK have relied upon rudimentary non-compartmental analyses with sparse time sampling which are not capable of addressing the complexities of phage PK. Furthermore, our analytical approach was critical to being able to assess both phage-specific and size-dependent effects. Though size dependency was not estimated with statistical significance, this is likely due to only having three different covariate levels (63 nm, 227 nm, and 400 nm). Additionally, this study confirms the ongoing utility of radiolabeling approaches to quantifying tissue concentrations of phages. Moving forward, nuclear medicine approaches will likely serve as an important tool for evaluating phage absorption, distribution, catabolism, and elimination in the absence of infection to provide a baseline for “ground truth” in phage disposition and elimination characteristics without the confounding effects of phage self-replication.

Evaluating absorption of phages, especially after oral, intraurethral, or inhalational administration is an outstanding question in the field of phage therapy. There are unclear mechanisms of phage transport across cellular barriers, which is underscored by previous studies which have found measurable phage concentrations in blood after oral or intraurethral administration(21, 23). Historically, oral bioavailability of biologics and biotic agents is assumed to be negligible due to the combined effects of drug due to low pH, digestive enzymes, and low intestinal wall permeability. Despite this longstanding PK assumption, multiple phage studies have highlighted the need to re-evaluate this assumption in the context of phage therapy(21, 23).

The final PBPK model describes phage distribution to each tissue using permeability and partition coefficients, which empirically account for phage tissue penetration. As a first-generation PBPK model of phages, this is a necessary first approach. However, transport of phages into the tissue is likely mediated by a combination of convective uptake (passive, size-dependent process) and active transport by endothelial cells(24). Recent work by Bichet and colleagues probed the question of transport of phage across cellular barriers and interestingly found a preferential transport direction of *apical-to-basolateral* sides(18). Thus, future studies should probe general phage transport rate and capacity across endothelial cells, as well as potential phage-determining effects (e.g., phage size or morphology).

Phage elimination was characterized using two predominant routes. First, elimination by the reticuloendothelial system is broadly considered the primary elimination route for particle-like drugs, and is well supported for liposomes, nanoparticles, and phages. However, our data and data from other groups has routinely found intravenously administered phages are capable of penetrating into the urine and gastric lumen. In the case of IV administered phage reaching the urine, penetration of the urine would be a physiological challenge owed to the 10 nm filtration cut- off, which is far smaller than the vast majority of phages being studied(25). Separately, orally administered phages must pass the intestinal wall, which is highly selective against microorganisms. In addition, immune-mediated clearance of phage, though outside the scope of this study, is another critical pathway of phage elimination. For acute infections, phage elimination by the innate immune system is likely most critical, as primary and secondary antibody response times are significantly longer (5-12 days) than the critical treatment period where phage is administered (i.e., ideally within 24h of onset, per clinical sepsis guidelines). However, for the types of chronic infections where phage therapy has been used under compassionate use circumstances, humoral immune response will likely be a significant route of clearance, too.

Radionuclide labeling of pharmaceuticals is a longstanding strategy to quantify PK. We chose to use the Sulfo-SHPP bi-functional conjugate for I-125 labeling of phage capsid proteins as it has the advantages of; 1) a protected aromatic ring to which oxidized I-125 iodide ion is bound, 2) an NHS ester which under physiologic conditions readily forms stable peptide (amide, covalent) bonds with accessible terminal amines of a protein and, 3) is water soluble at a neutral pH. We also chose a two-step method for phage radiolabeling by first oxidatively attaching I-125 to Sulfo- SHPP™ using Iodobeads™ which were then removed before adding to phage for a brief conjugation reaction. This two-step approach prevents oxidative damage and over modification of phage to preserve structural integrity and infectivity. I-125-SHPP-labeled phage is highly resistant to spontaneous or enzymatic dehalogenation or proteolysis in vivo and ideal for PK/PD studies in pre-clinical models.

Moving forward, PK studies of phage should focus on evaluating the effects of phage size and structure on distribution and elimination, but other salient features of phage biology should also be evaluated. For example, nanoparticles for pharmaceutical development are often characterized for their zeta potential, which has been shown to influence both formulation stability and pharmacokinetics. Optimal zeta potential has been reported as being between -30 and +30, which coincides with previous reports of phage’s anionic zeta potential. Ultimately, the future of phage therapy as an antibiotic class orthogonal to small molecule antibiotics is incredibly promising in the context of contemporary drug development strategies used for other biotic agents. If realized, phage therapy could restore the antimicrobial armamentarium and provide clinicians with a valuable resource in treating highly drug-resistant bacterial pathogens.

## Acknowledgements

Research reported in this publication was supported by the National Institute of Allergy and Infectious Diseases of the National Institutes of Health under Award Number R01AI177997 (**PD/PI Smith**). The content is solely the responsibility of the authors and does not necessarily represent the official views of the National Institutes of Health.

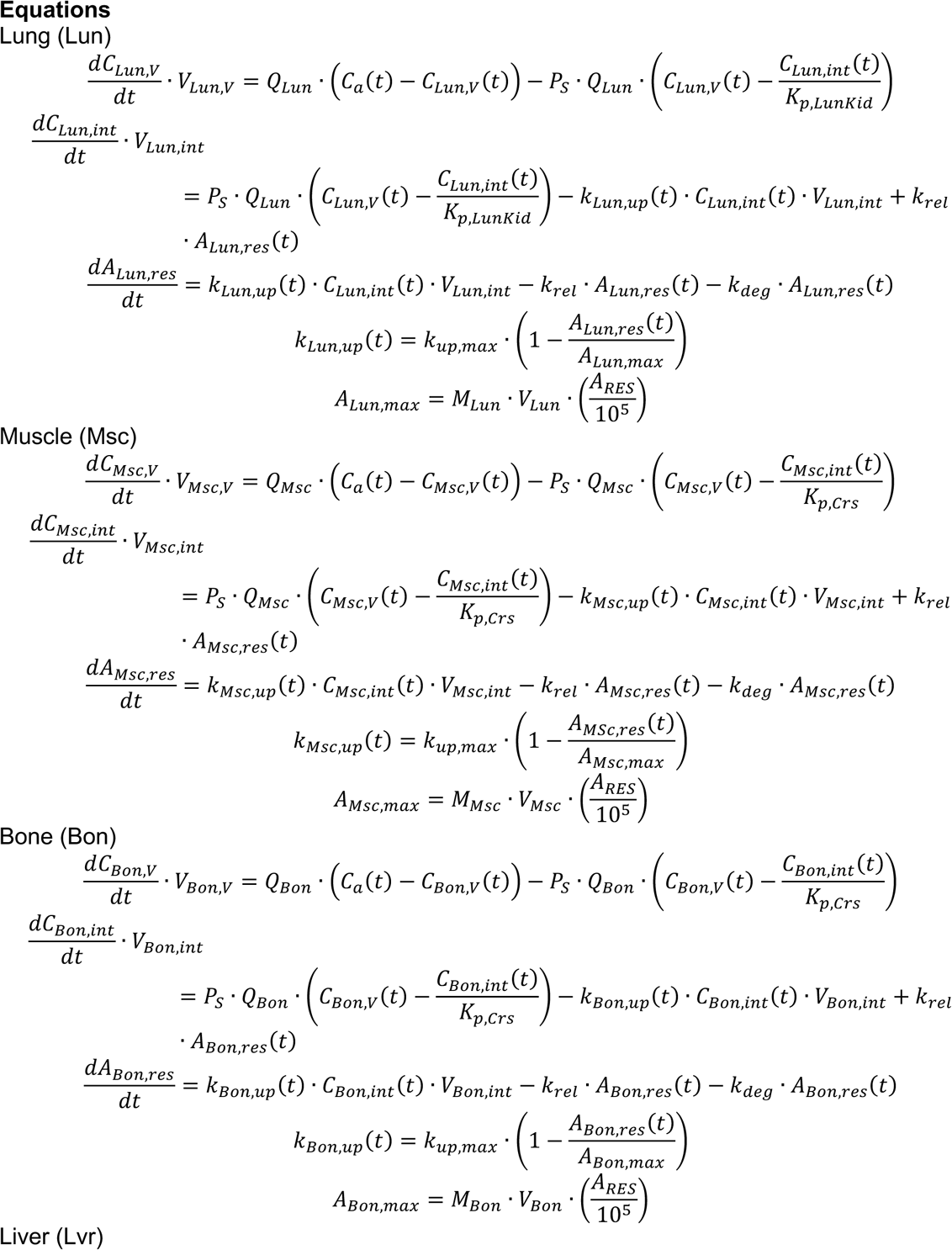

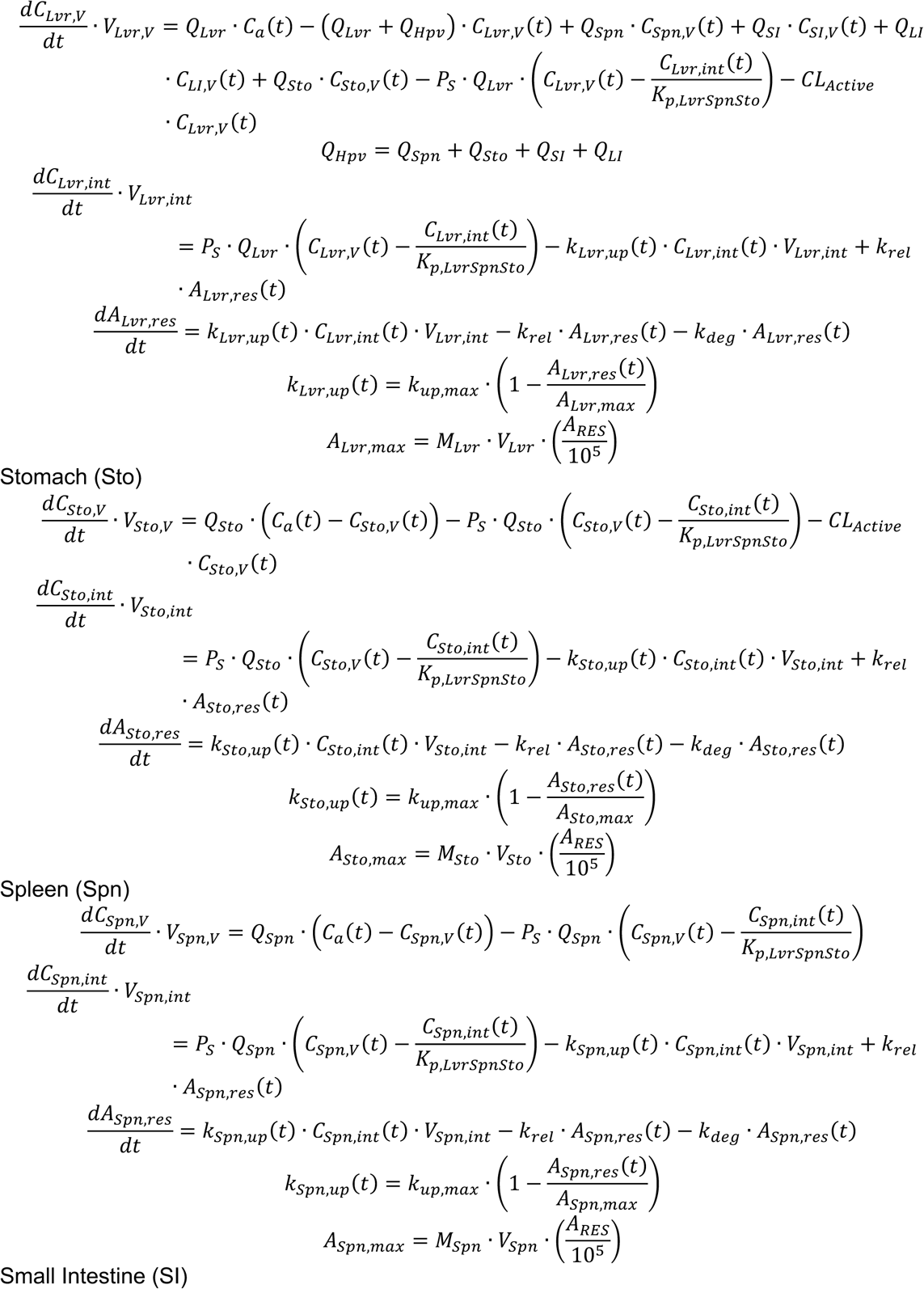

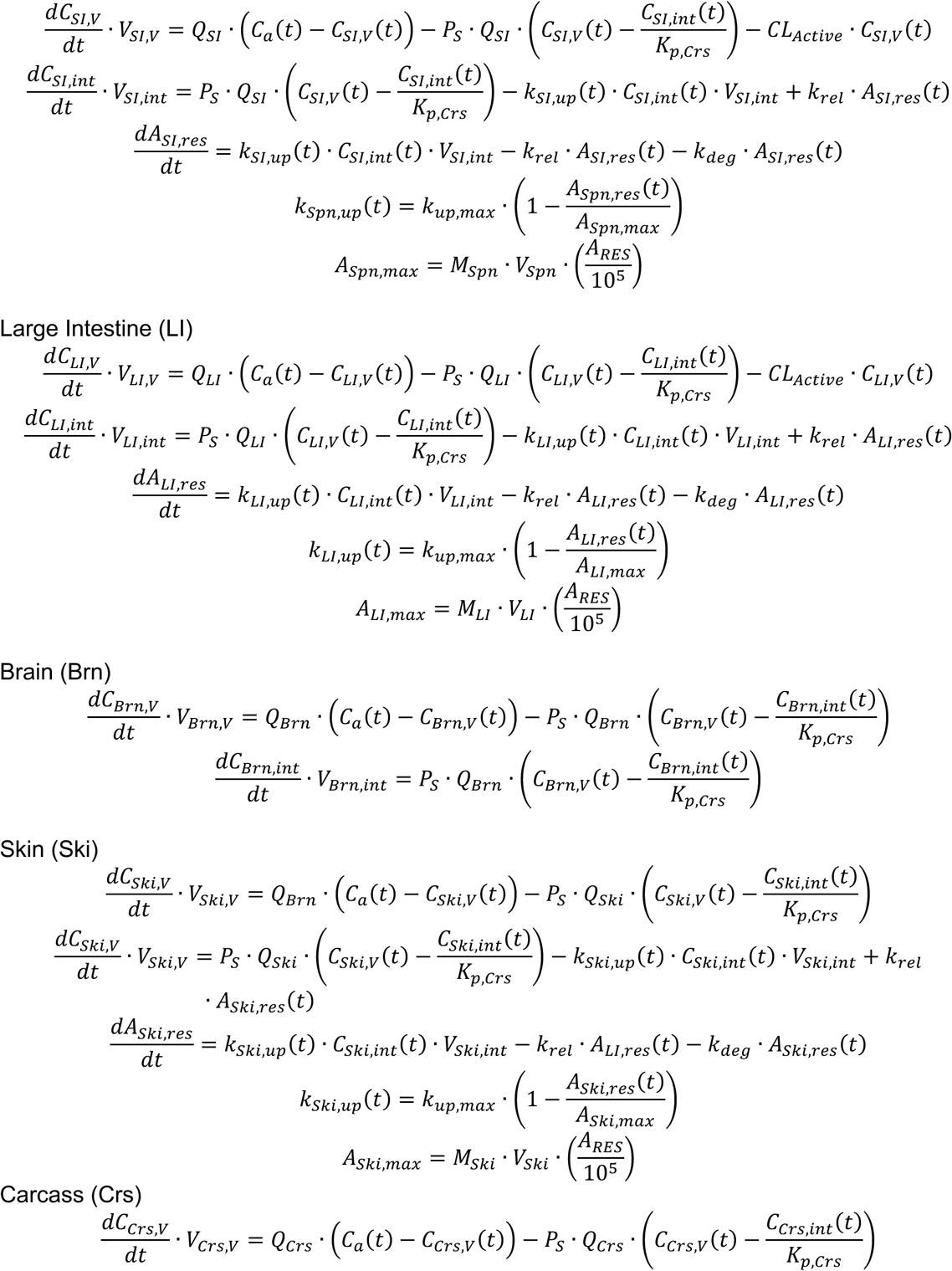

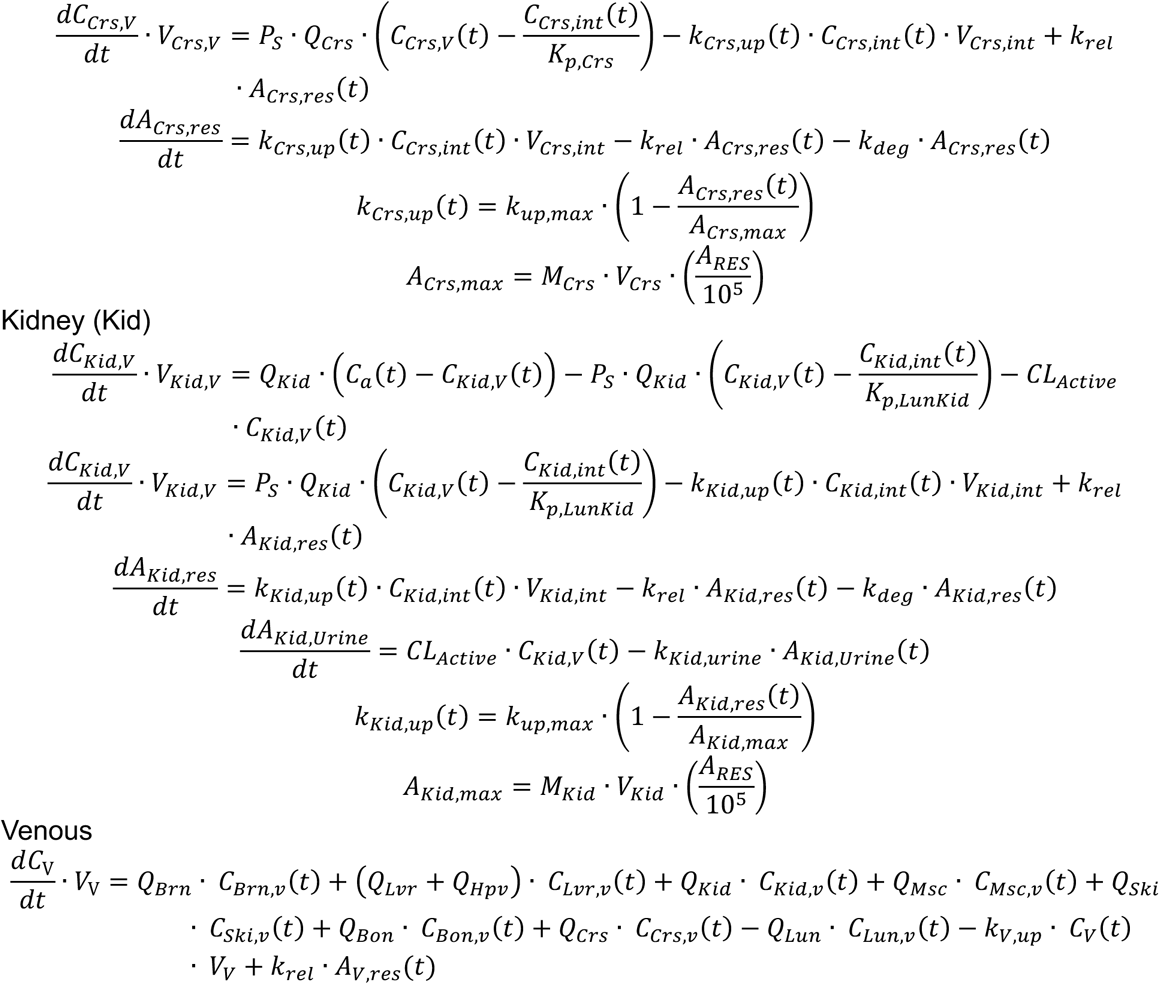

## SUPPLEMENTAL MATERIALS

**Figure S1:**
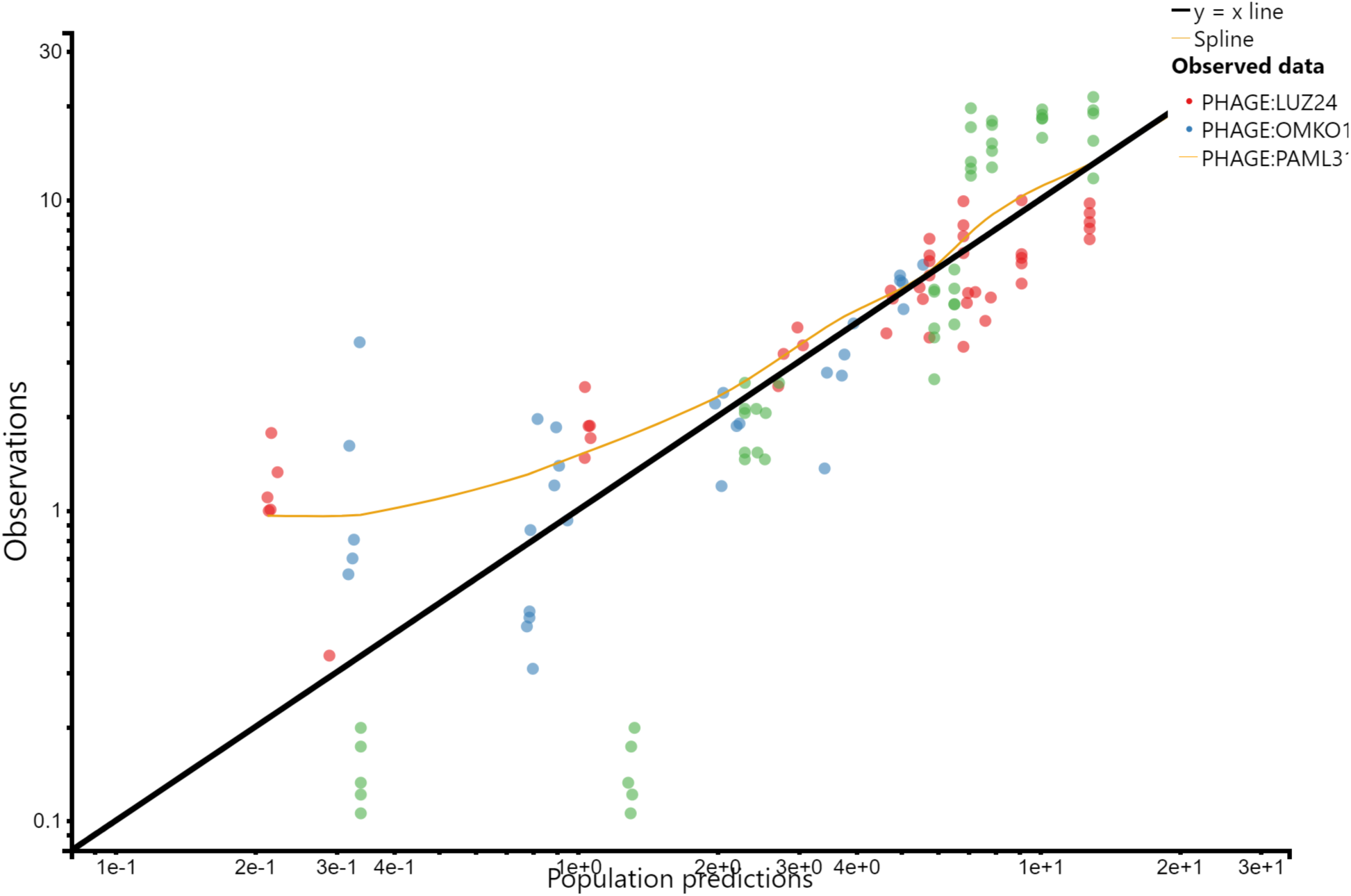
Observed versus predicted diagnostic plot of Blood measurements.

**Figure S2:**
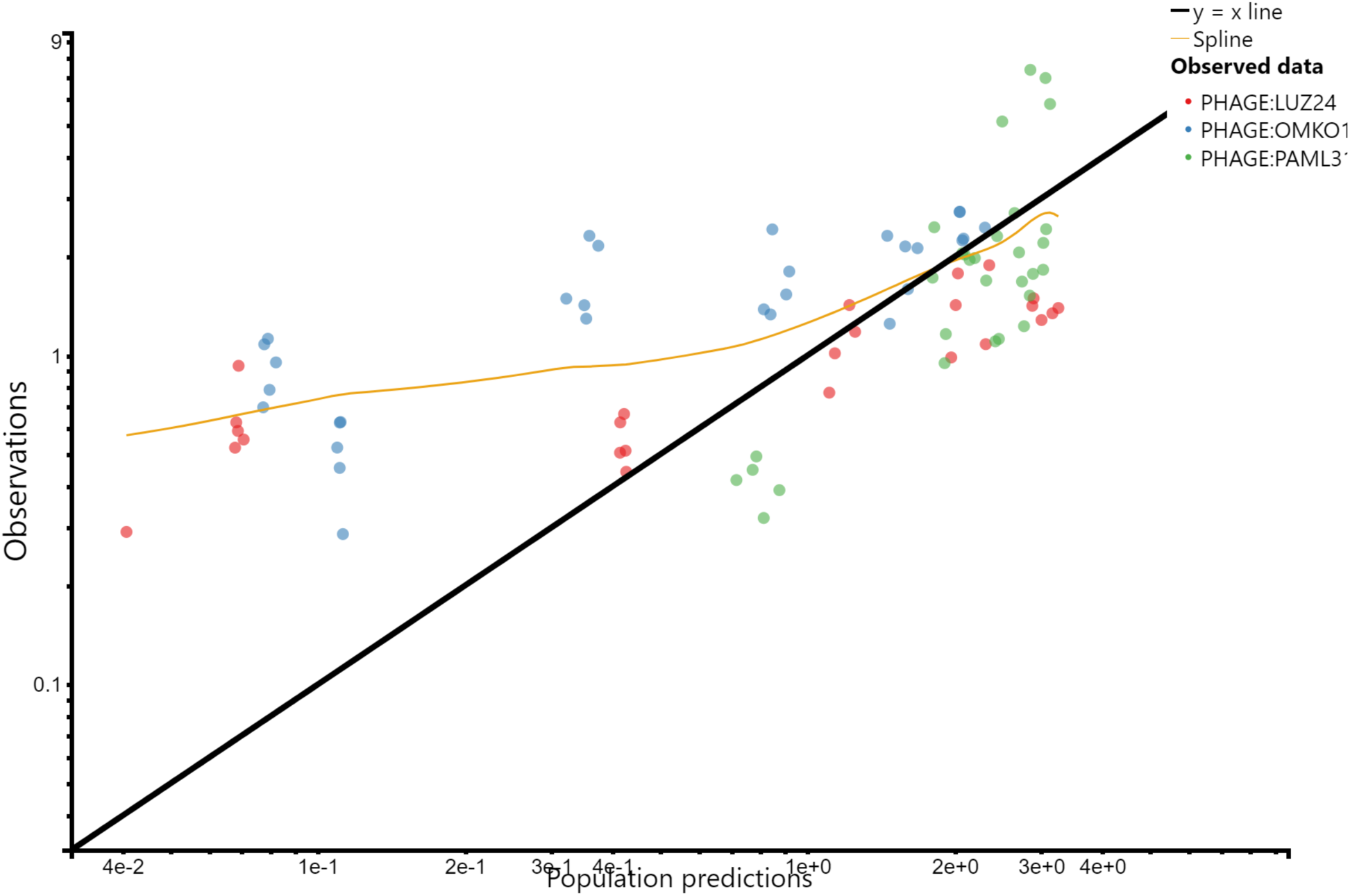
Observed versus predicted diagnostic plot of Bone measurements.

**Figure S3:**
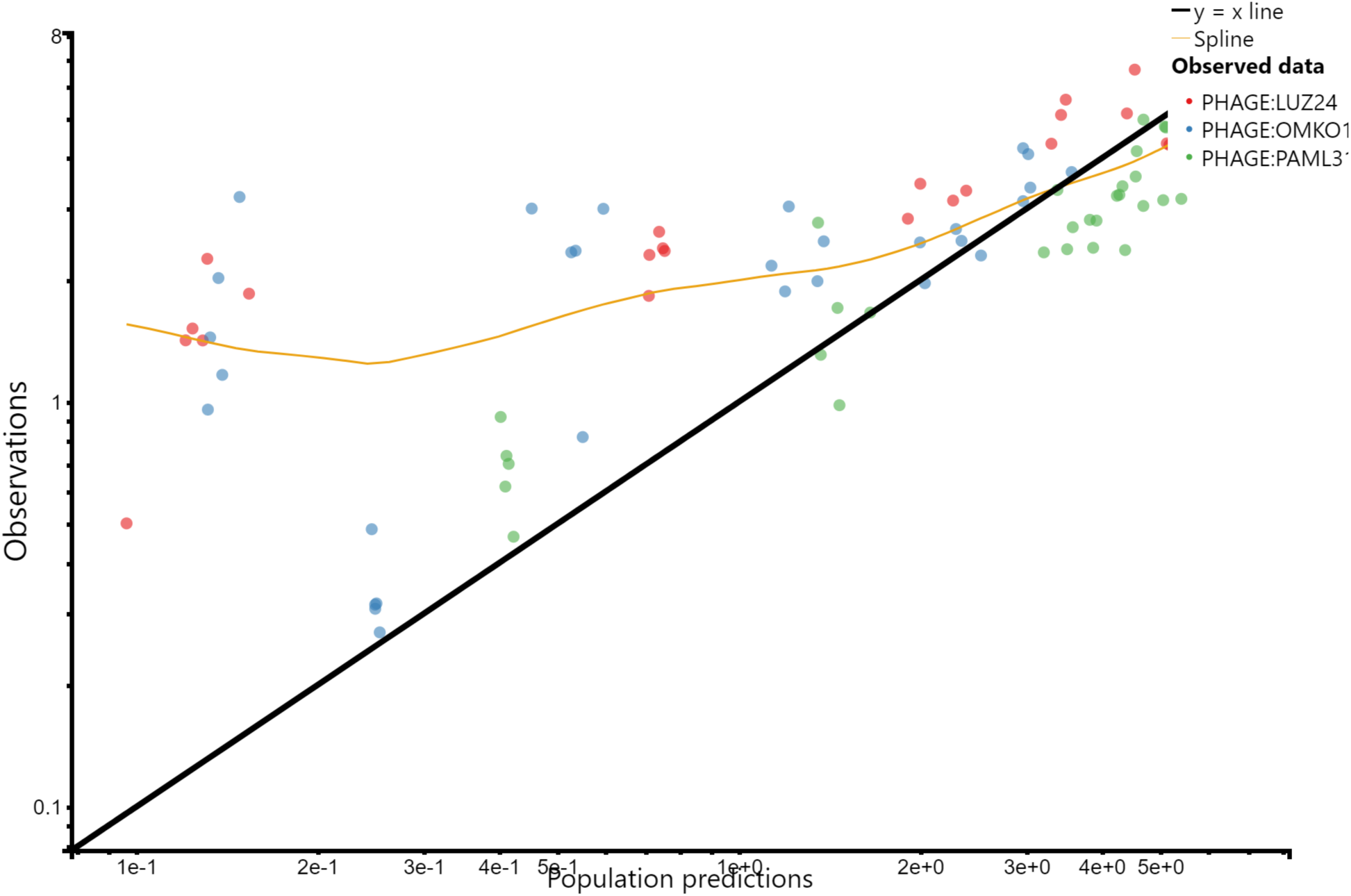
Observed versus predicted diagnostic plot of Kidney measurements.

**Figure S4:**
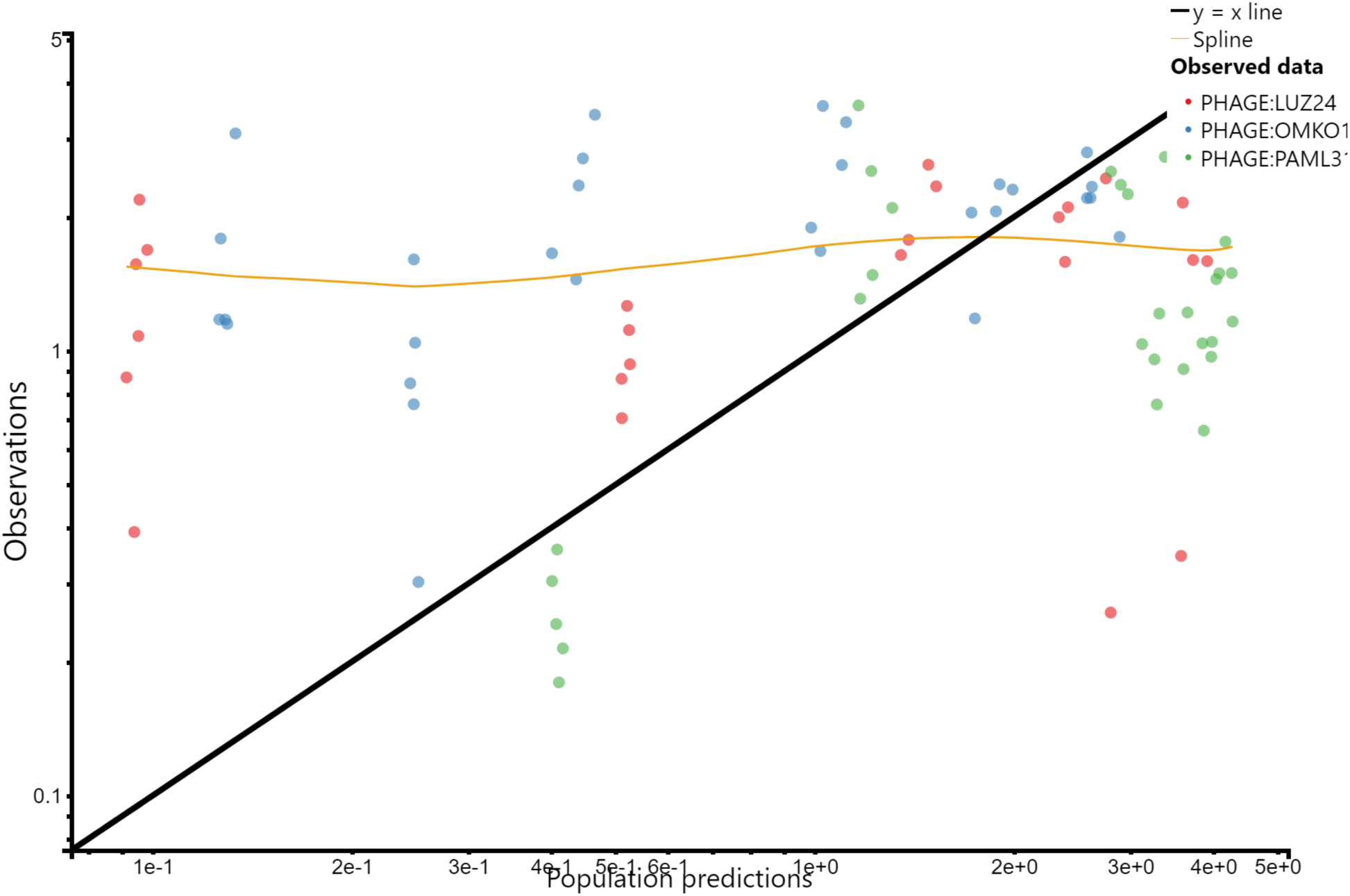
Observed versus predicted diagnostic plot of Lg. Intestines measurements.

**Figure S5:**
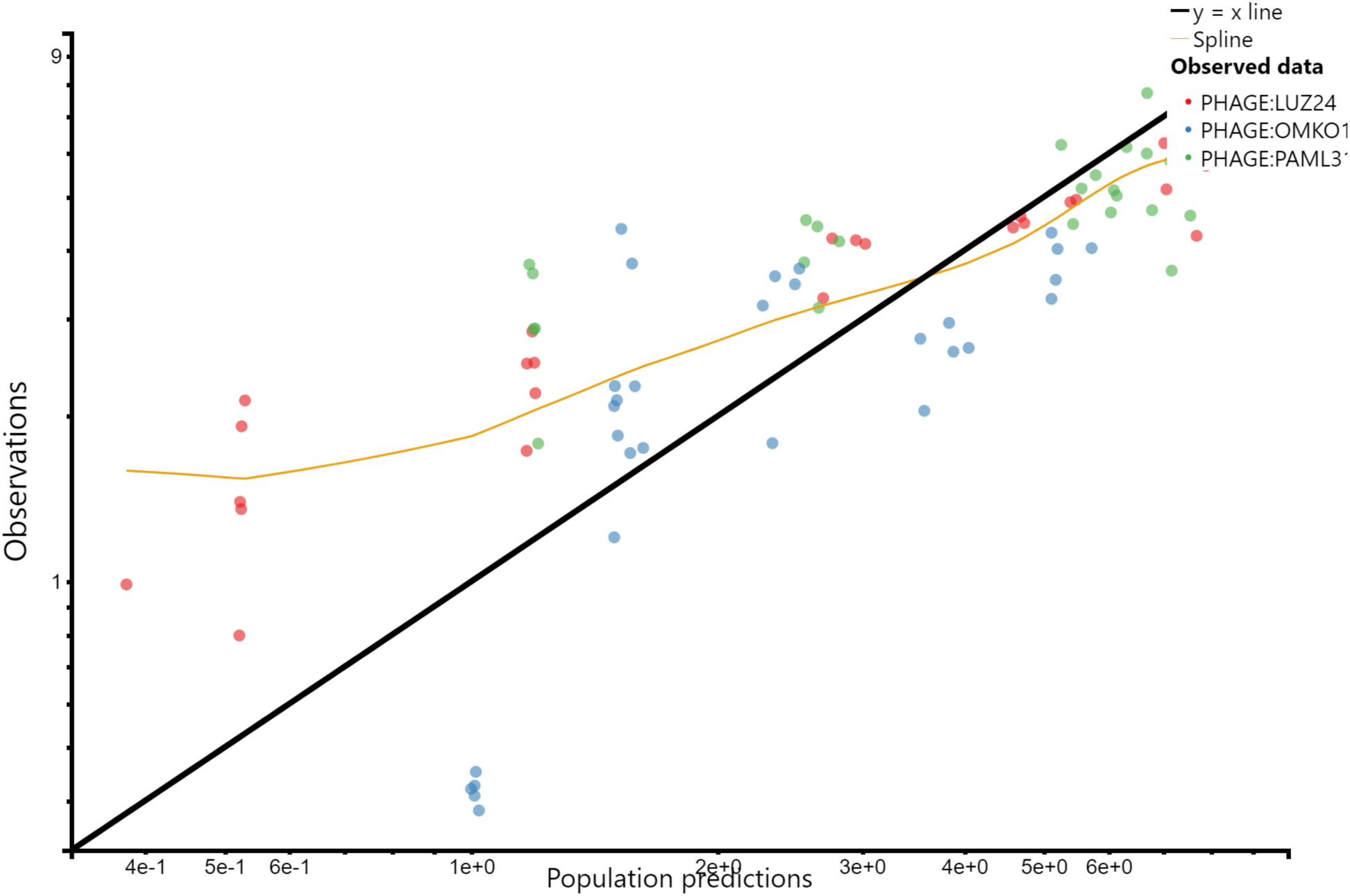
Observed versus predicted diagnostic plot of Liver measurements.

**Figure S6:**
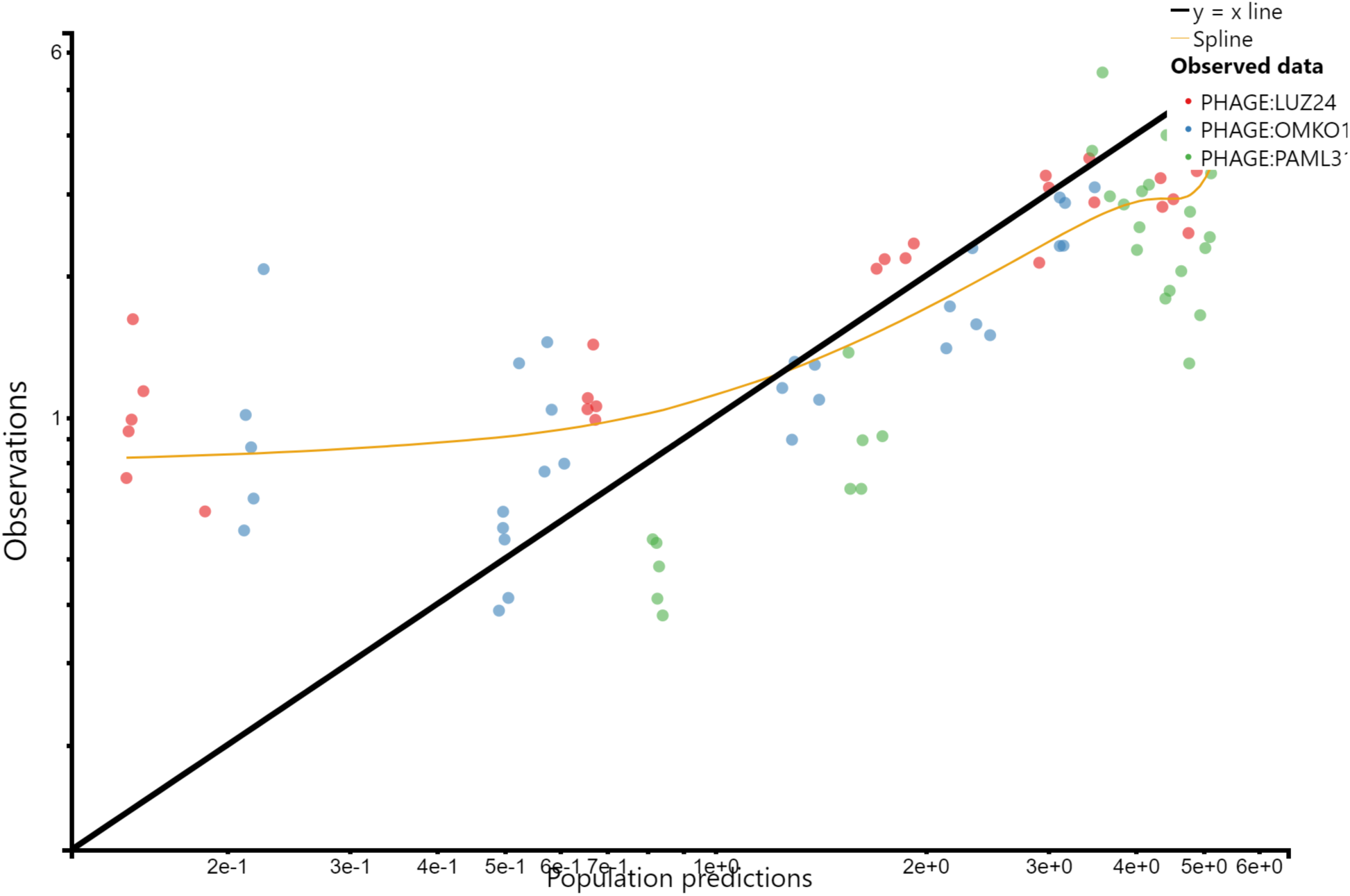
Observed versus predicted diagnostic plot of Lung measurements.

**Figure S7:**
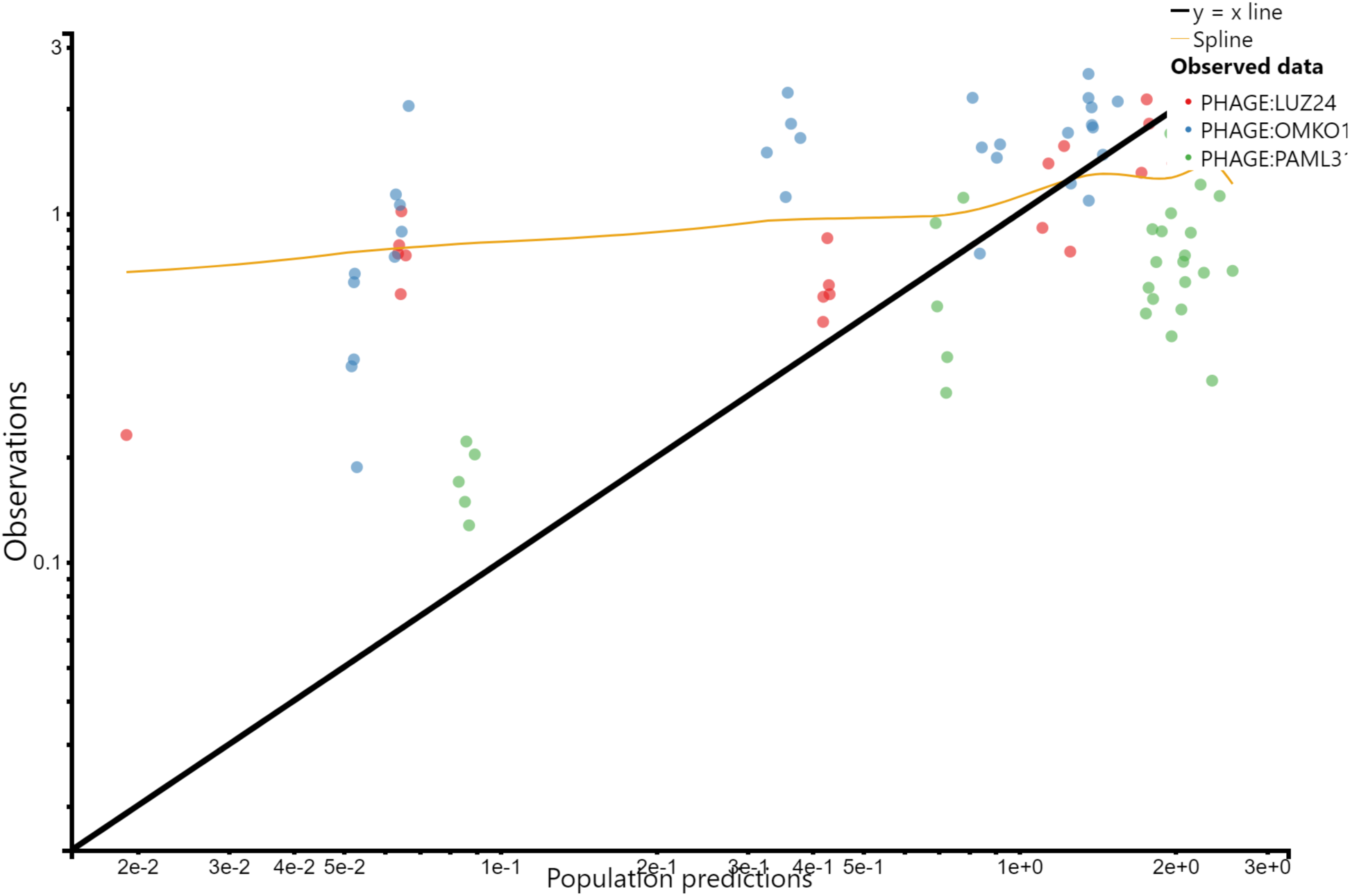
Observed versus predicted diagnostic plot of Muscle measurements.

**Figure S8:**
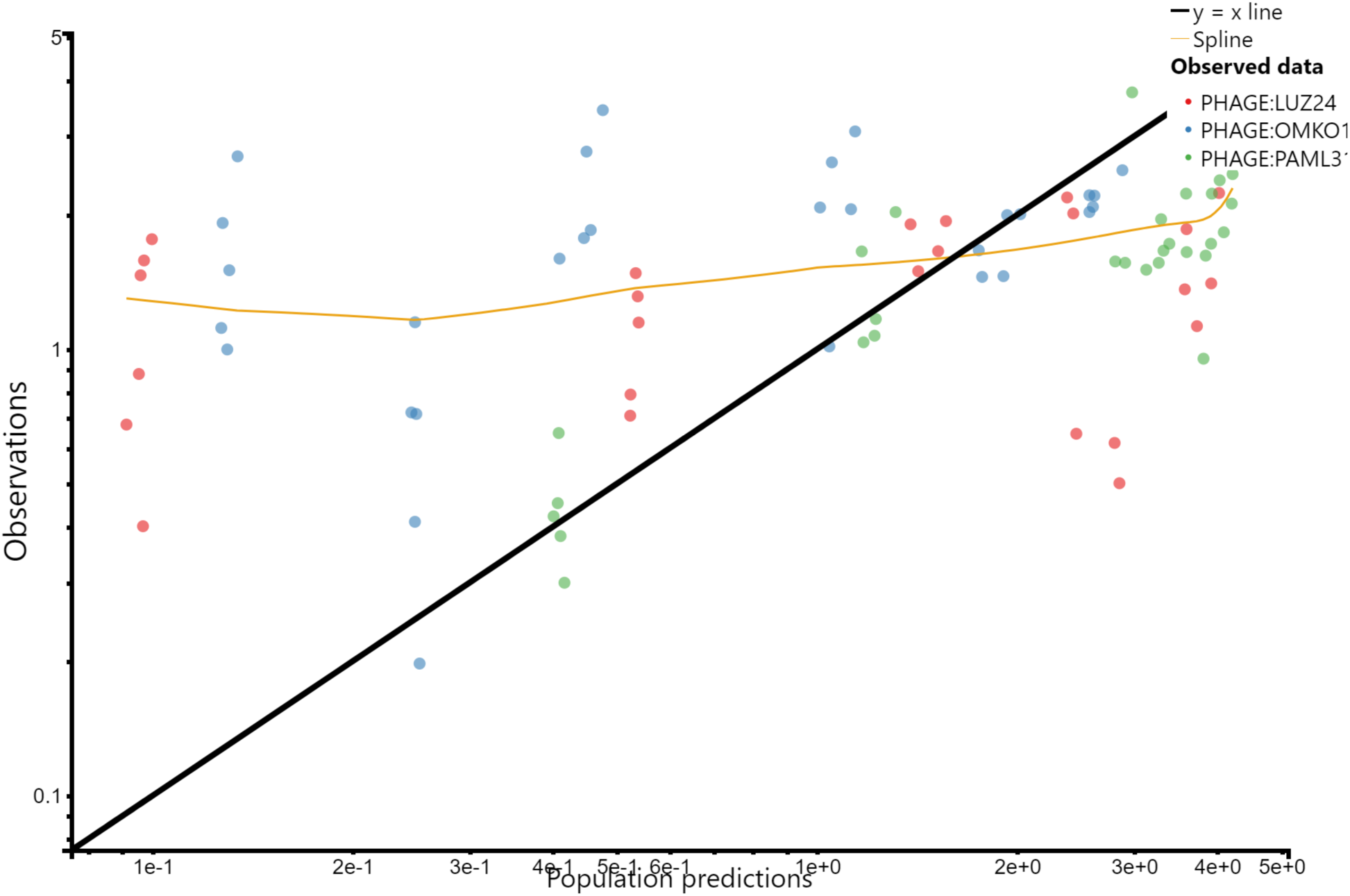
Observed versus predicted diagnostic plot of Sm. Intestines measurements.

**Figure S9:**
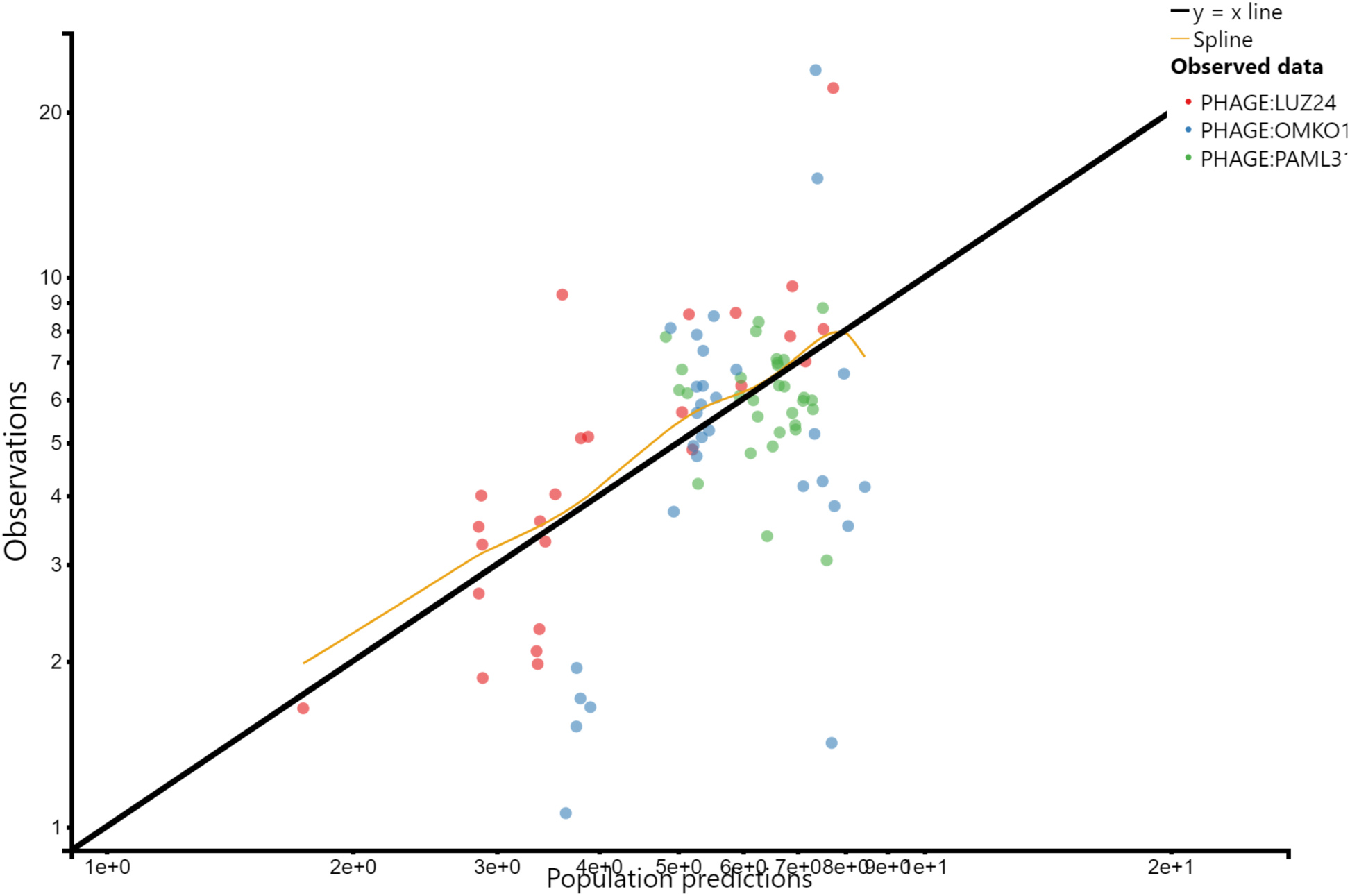
Observed versus predicted diagnostic plot of Spleen measurements.

**Figure S10:**
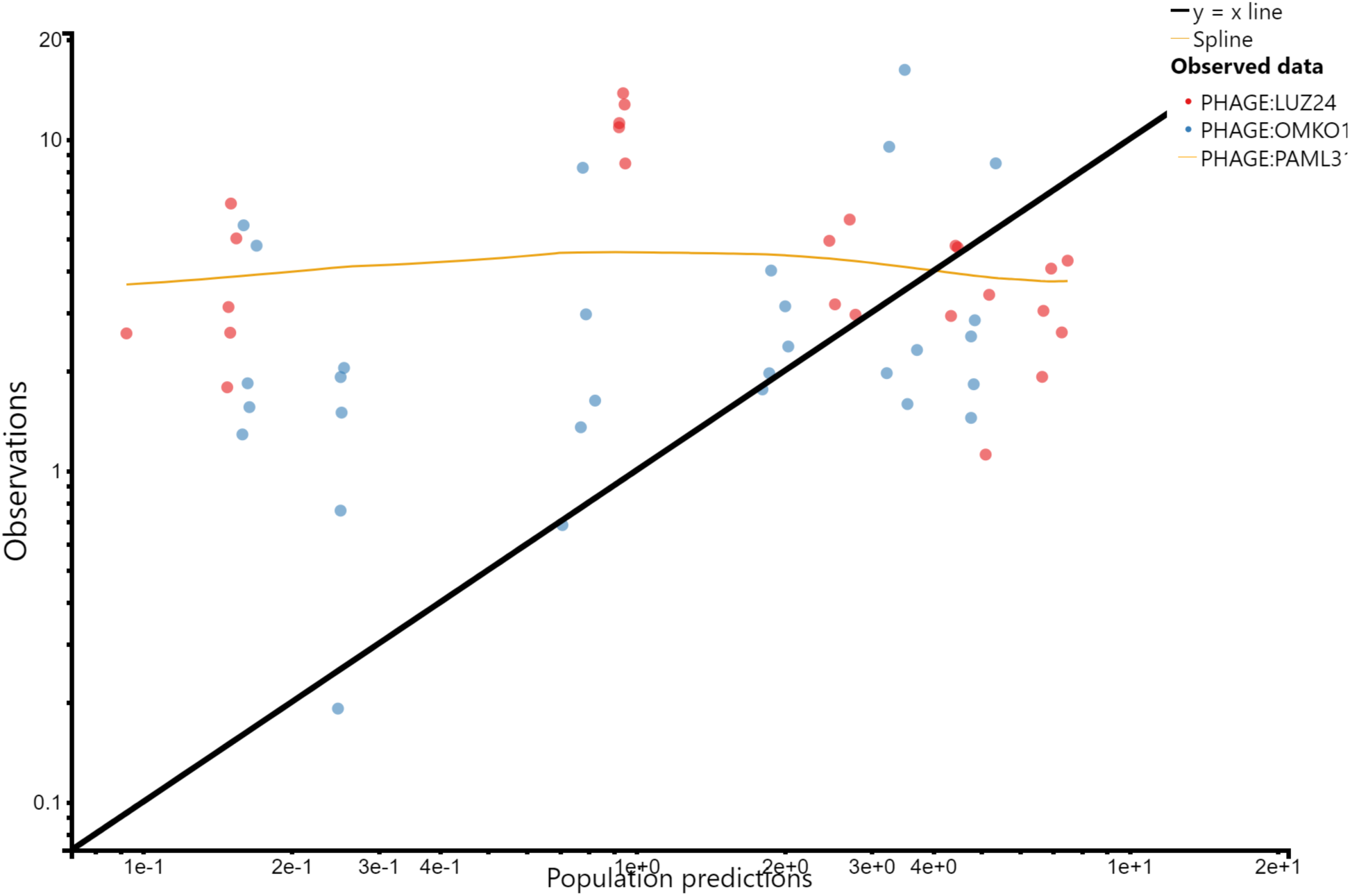
Observed versus predicted diagnostic plot of Stomach measurements.

**Figure S11:**
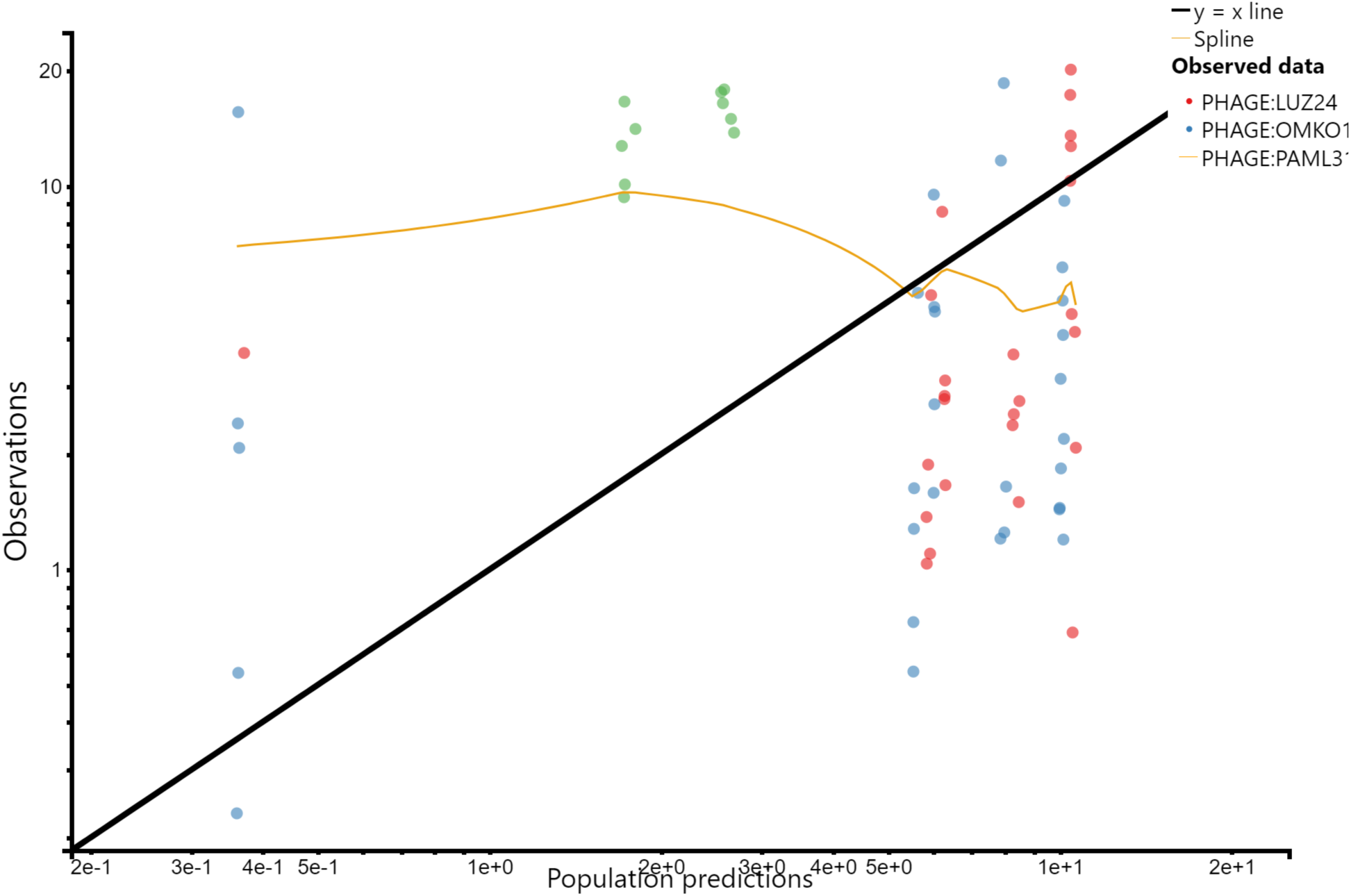
Observed versus predicted diagnostic plot of Stomach Contents measurements.

**Figure S12:**
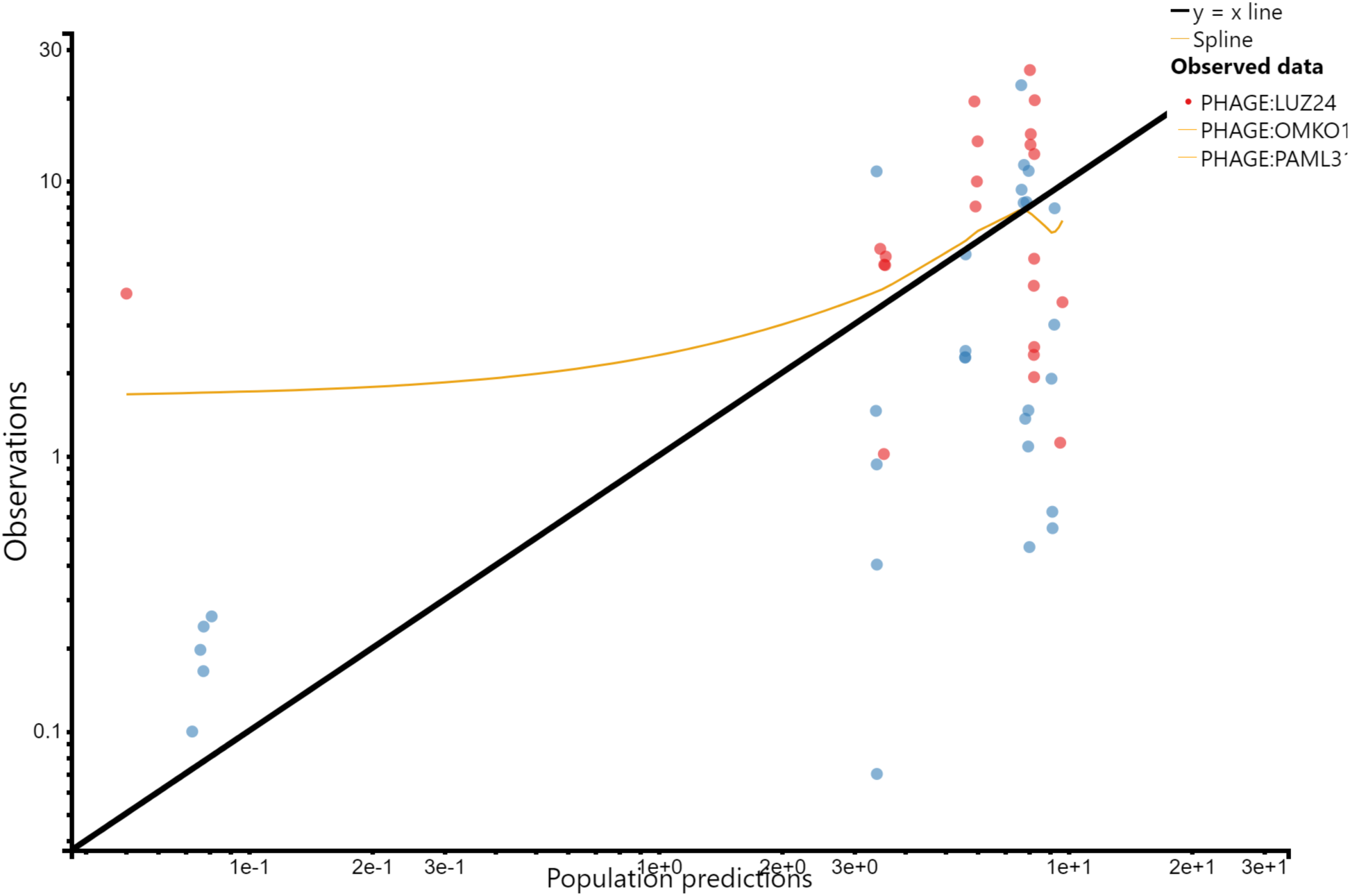
Observed versus predicted diagnostic plot of Urine measurements.

**Figure S13:**
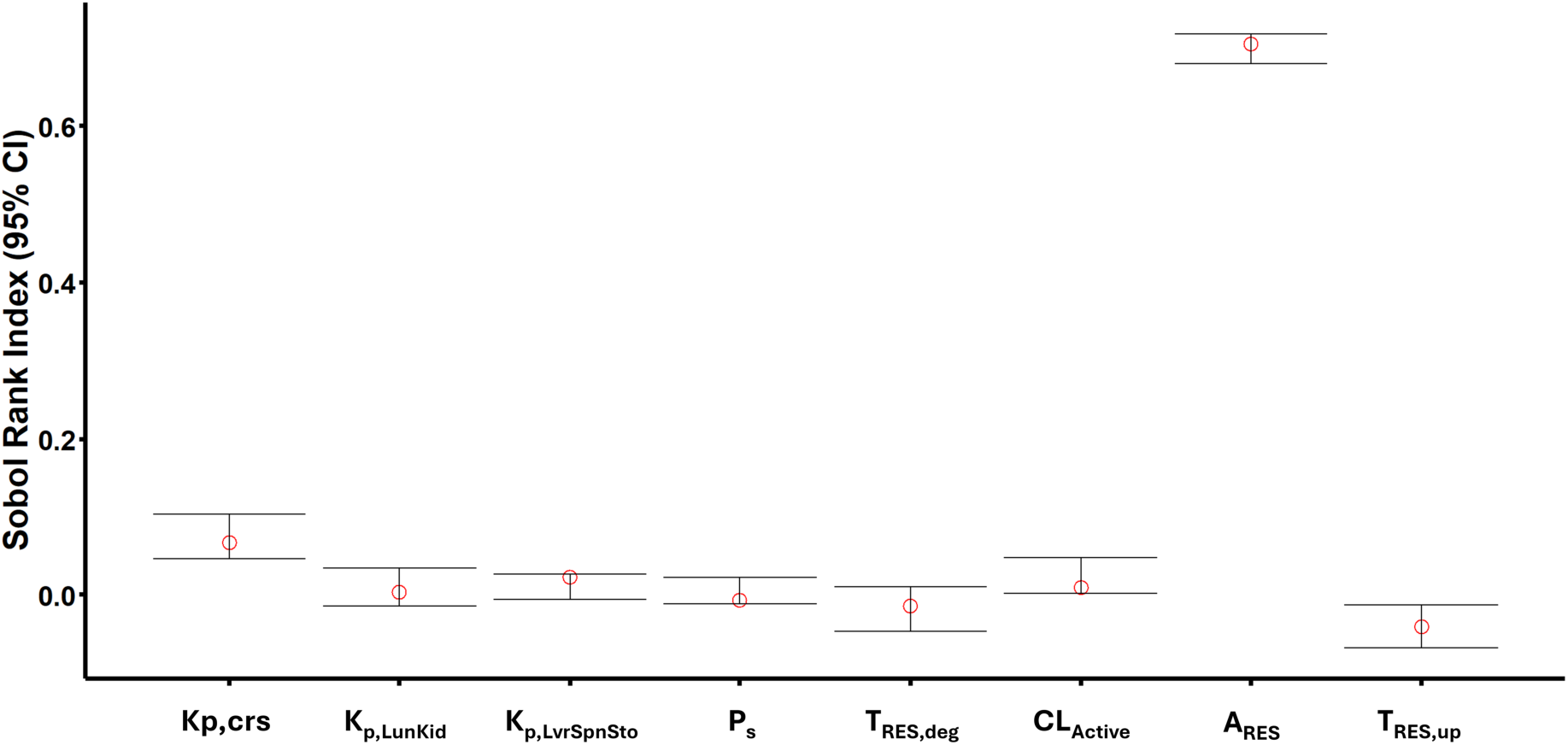
Global Sensitivity Analysis (GSA) of key model parameters

**Figure S14:**
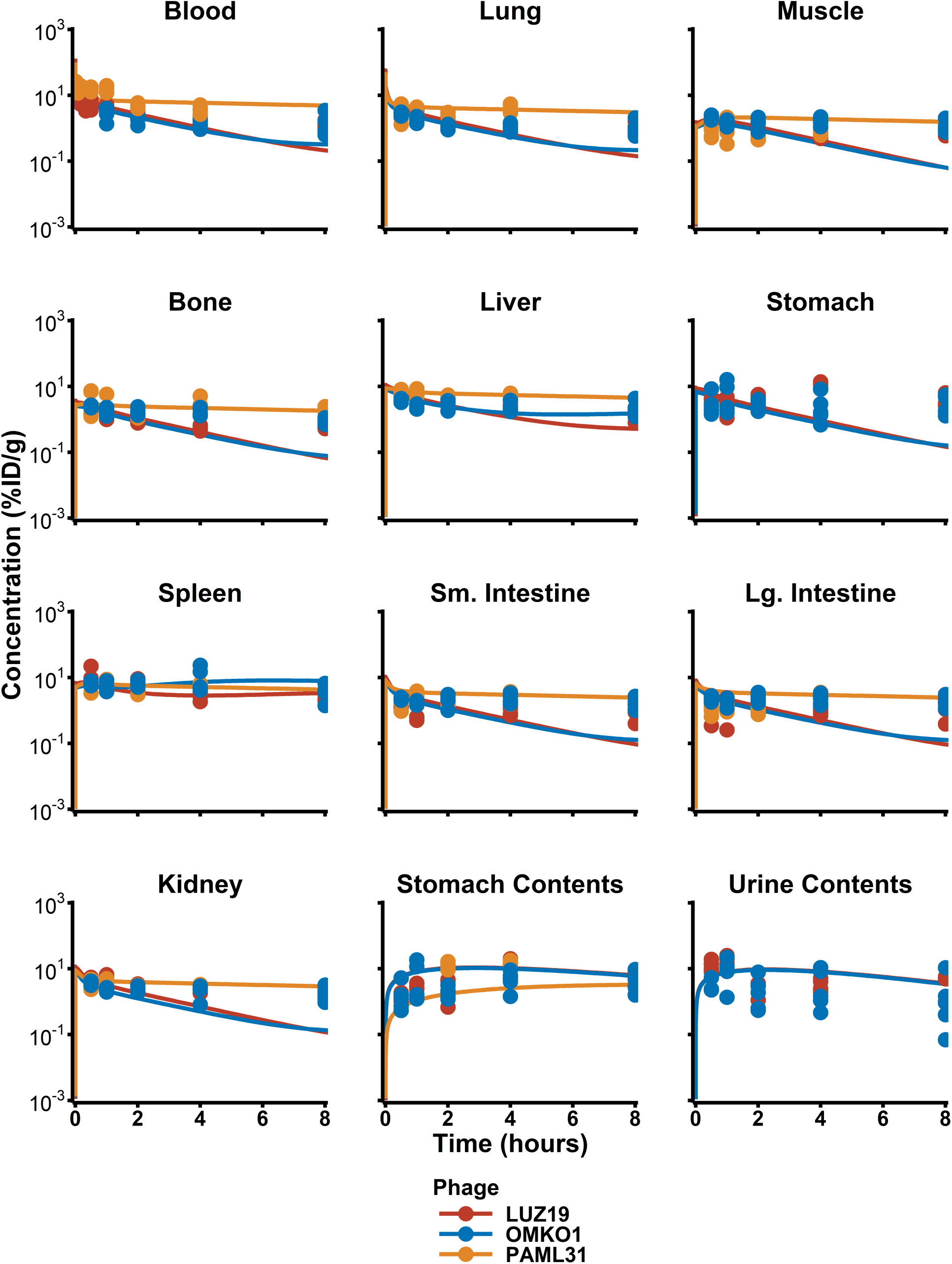
**PBPK Post Hoc fits, first 8 hours**

**Table S1:**
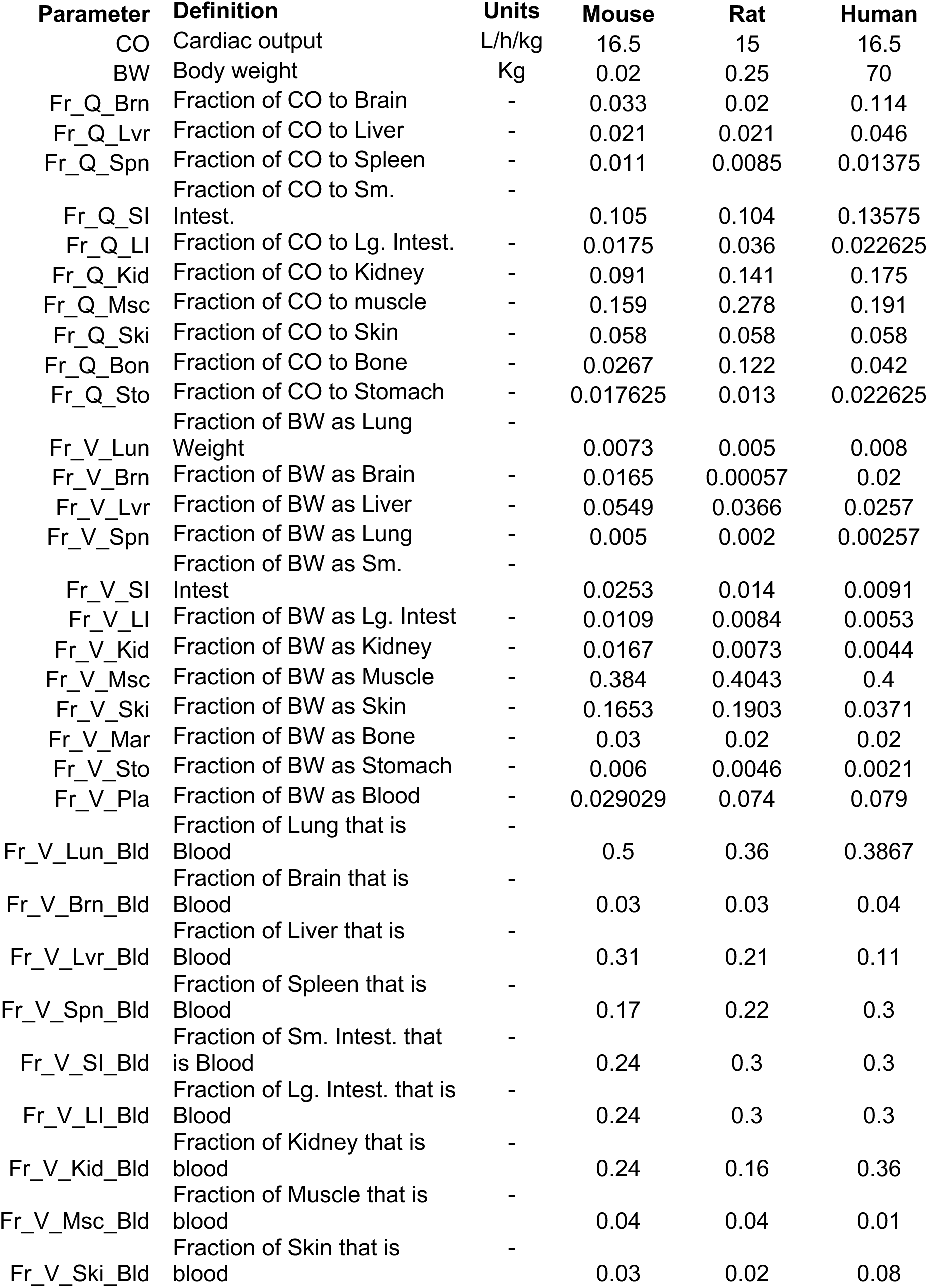
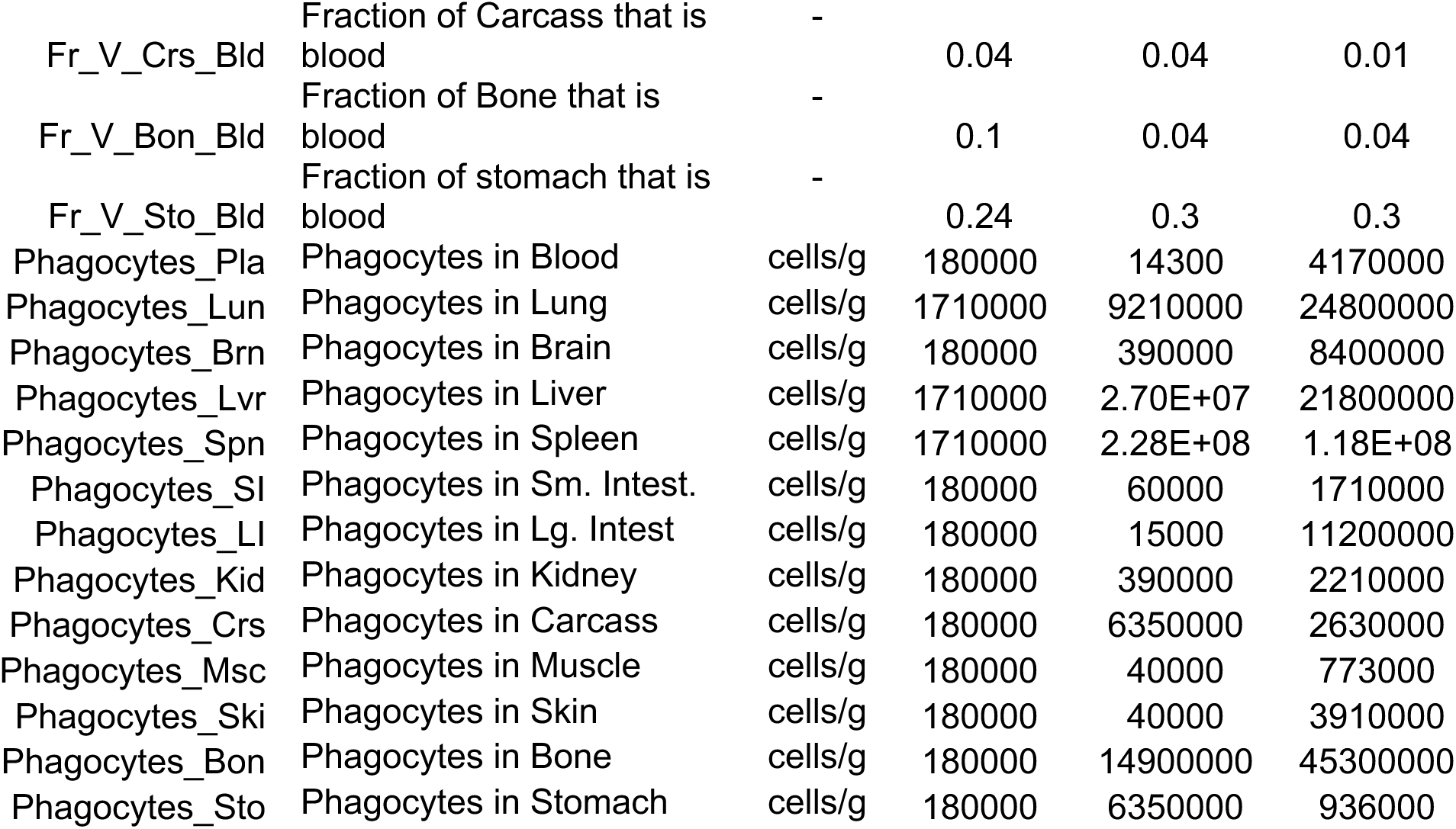
PBPK Model Parameters.

**Table S2:** Global Sensitivity Analysis (GSA) of Key Model Parameters. First order rank Sobol indices for each parameter based on 1000 simulated parameter sets and 95% confidence intervals.

|  | First-order rank | bias | std. error | min. c.i. | max. c.i. |
| --- | --- | --- | --- | --- | --- |
| $K_{p,Crs}$ | 0.066631 | -0.00678 | 0.020091 | 0.045669 | 0.102709 |
| $K_{p, LunKid}$ | 0.003565 | -0.00394 | 0.016095 | -0.01503 | 0.034368 |
| $K_{p, LvrSpnSto}$ | 0.021637 | 0.006517 | 0.010788 | -0.00516 | 0.026982 |
| $P_S$ | -0.00742 | -0.01114 | 0.010871 | -0.01188 | 0.021841 |
| $T_{MPS, Deg}$ | -0.01457 | 0.003462 | 0.019097 | -0.04674 | 0.009832 |
| $CL_{Active}$ | 0.009487 | -0.00908 | 0.016183 | 0.002325 | 0.046665 |
| $A_{MPS}$ | 0.704755 | 0.001264 | 0.010696 | 0.679477 | 0.719241 |
| $T_{MPS, Up}$ | -0.04068 | 0.001177 | 0.01736 | -0.06836 | -0.01349 |

## References

1. CDC. In: US Department of Health and Human Services ed. Atlanta: CDC; 2019.

2. Tacconelli E, Carrara E, Savoldi A, Harbarth S, Mendelson M, Monnet DL, et al. Discovery, research, and development of new antibiotics: the WHO priority list of antibiotic-resistant bacteria and tuberculosis. The Lancet Infectious diseases. 2018;18(3):318–27.

3. O’neill J. Antimicrobial resistance: tackling a crisis for the health and wealth of nations. Rev Antimicrob Resist. 2014.

4. Kaye KS, and Pogue JM. Infections Caused by Resistant Gram-Negative Bacteria: Epidemiology and Management. Pharmacotherapy: The Journal of Human Pharmacology and Drug Therapy. 2015;35(10):949–62.

5. Denissen J, Reyneke B, Waso-Reyneke M, Havenga B, Barnard T, Khan S, et al. Prevalence of ESKAPE pathogens in the environment: Antibiotic resistance status, community-acquired infection and risk to human health. International Journal of Hygiene and Environmental Health. 2022;244:114006.

6. Montero MM, López Montesinos I, Knobel H, Molas E, Sorlí L, Siverio-Parés A, et al. Risk Factors for Mortality among Patients with Pseudomonas aeruginosa Bloodstream Infections: What Is the Influence of XDR Phenotype on Outcomes? J Clin Med. 2020;9(2).

7. Zhang Y, Chen XL, Huang AW, Liu SL, Liu WJ, Zhang N, et al. Mortality attributable to carbapenem-resistant Pseudomonas aeruginosa bacteremia: a meta-analysis of cohort studies. Emerging microbes & infections. 2016;5(3):e27.

8. Liu Q, Li X, Li W, Du X, He JQ, Tao C, et al. Influence of carbapenem resistance on mortality of patients with Pseudomonas aeruginosa infection: a meta-analysis. Sci Rep. 2015;5:11715.

9. D’Herelle F. Sur un microbe invisible antagoniste des bacilles dysentériques. CR Acad Sci. 1917;165:373–5.

10. Suh GA, Lodise TP, Tamma PD, Knisely JM, Alexander J, Aslam S, et al. Considerations for the Use of Phage Therapy in Clinical Practice. Antimicrobial agents and chemotherapy. 2022;66(3):e0207121.

11. Jia HJ, Jia PP, Yin S, Bu LK, Yang G, and Pei DS. Engineering bacteriophages for enhanced host range and efficacy: insights from bacteriophage-bacteria interactions. Frontiers in microbiology. 2023;14:1172635.

12. Yehl K, Lemire S, Yang AC, Ando H, Mimee M, Torres MT, et al. Engineering Phage Host-Range and Suppressing Bacterial Resistance through Phage Tail Fiber Mutagenesis. Cell. 2019;179(2):459–69.e9.

13. Wang H, Yang Y, Xu Y, Chen Y, Zhang W, Liu T, et al. Phage-based delivery systems: engineering, applications, and challenges in nanomedicines. Journal of Nanobiotechnology. 2024;22(1):365.

14. Smith NM, Nguyen TD, Chin WH, Sanborn JT, Souza Hd, Ho BM, et al. A Mechanism-based Pathway Toward Administering Highly Active N-Phage Cocktails. Frontiers Microbiology. 2023.

15. Siopi M, Skliros D, Paranos P, Koumasi N, Flemetakis E, Pournaras S, et al. Pharmacokinetics and pharmacodynamics of bacteriophage therapy: a review with a focus on multidrug-resistant Gram-negative bacterial infections. Clinical Microbiology Reviews. 2024;37(3):e00044–24.

16. Zhang M, Gao S, Yang D, Fang Y, Lin X, Jin X, et al. Influencing factors and strategies of enhancing nanoparticles into tumors in vivo. Acta Pharm Sin B. 2021;11(8):2265–85.

17. Echterhof A, Dharmaraj T, Khosravi A, McBride R, Miesel L, Chia J-H, et al. The contribution of neutrophils to bacteriophage clearance and pharmacokinetics in vivo. JCI Insight. 2024;9(20).

18. Bichet MC, Chin WH, Richards W, Lin Y-W, Avellaneda-Franco L, Hernandez CA, et al. Bacteriophage uptake by mammalian cell layers represents a potential sink that may impact phage therapy. iScience. 2021;24(4):102287.

19. Iooss B. online: CRAN; 2024.

20. Lin YW, Chang RY, Rao GG, Jermain B, Han ML, Zhao JX, et al. Pharmacokinetics/pharmacodynamics of antipseudomonal bacteriophage therapy in rats: a proof-of-concept study. Clinical Microbiology and Infection. 2020;26(9):1229–35.

21. Kim P, Sanchez AM, Penke TJR, Tuson HH, Kime JC, McKee RW, et al. Safety, pharmacokinetics, and pharmacodynamics of LBP-EC01, a CRISPR-Cas3-enhanced bacteriophage cocktail, in uncomplicated urinary tract infections due to *Escherichia coli* (ELIMINATE): the randomised, open-label, first part of a two-part phase 2 trial. *The Lancet Infectious Diseases*.

22. Nguyen TD, Smith NM, Attwood K, Gundroo A, Chang S, Yonis M, et al. Bayesian optimization of tacrolimus exposure in stable kidney transplant patients. Pharmacotherapy: The Journal of Human Pharmacology and Drug Therapy. 2023;43(10):1032–42.

23. Dhungana G, Nepal R, Regmi M, and Malla R. Pharmacokinetics and Pharmacodynamics of a Novel Virulent Klebsiella Phage Kp_Pokalde_002 in a Mouse Model. Frontiers in Cellular and Infection Microbiology. 2021;11.

24. Chen B, Dong JQ, Pan WJ, and Ruiz A. Pharmacokinetics/pharmacodynamics model- supported early drug development. Curr Pharm Biotechnol. 2012;13(7):1360–75.

25. Du B, Jiang X, Das A, Zhou Q, Yu M, Jin R, et al. Glomerular barrier behaves as an atomically precise bandpass filter in a sub-nanometre regime. Nat Nanotechnol. 2017;12(11):1096–102.

